# Profiling of terminating ribosomes reveals translational control at stop codons

**DOI:** 10.1101/2025.09.16.676599

**Authors:** Longfei Jia, Yuanhui Mao, Saori Uematsu, Xinyi Ashley Liu, Leiming Dong, Leonardo Henrique França de Lima, Shu-Bing Qian

**Affiliations:** Division of Nutritional Sciences, Cornell University, Ithaca, NY 14853, USA; Laboratory of Molecular Modelling and Bioinformatics (LAMMB), Department of Physical and Biological Sciences, Campus Sete Lagoas, Universidade Federal de São João Del Rei, Sete Lagoas, Brazil; State Key Laboratory of Reproductive Medicine and Offspring Health, Nanjing Medical University, Nanjing, P.R.China; Liangzhu Laboratory, Zhejiang University, Hangzhou, P.R.China

## Abstract

Accurate termination of protein synthesis is paramount for the integrity of cellular proteome, yet the dynamics and fidelity of ribosome termination remain poorly understood. Here, we establish a profiling strategy to capture terminating ribosomes in mammalian cells and reveal a substantial heterogeneity in ribosome pausing at individual stop codons. We identify a sequence motif upstream of the stop codon that promotes termination pausing, a finding validated by massively paralleled reporter assays. Unexpectedly, reduced termination pausing increases the likelihood of stop codon slippage, giving rise to proteins with heterogenous C-terminal extensions. Mechanistically, we show that sequence-dependent termination pausing arises from post-decoding mRNA scanning by the 3’ end of 18S rRNA. We further uncover tissue-specific patterns of termination pausing that correlates with the stoichiometry of Rps26, which modulates mRNA:rRNA interactions. Together, these results establish termination pausing as a distinct translational signature shaped by mRNA sequence contexts, ribosome heterogeneity, and cell type-specific translational control.

## INTRODUCTION

Eukaryotic mRNA translation terminates when a stop codon (UAA, UAG, or UGA) enters the ribosomal A site, triggering the coordinated actions of the release factors eRF1 and eRF3 ^1^. Structurally mimicking a tRNA, eRF1 mediates stop codon recognition and catalyzes hydrolysis of the peptidyl-tRNA bond, while eRF3 promotes this process in a GTP-dependent manner ^2^. In parallel, the ribosome recycling factor ABCE1 facilitates completion of termination by promoting dissociation of the post-termination ribosome ^3,4^. Following Upon release of the nascent polypeptide from the P-site, ABCE1 drives splitting of the 80S ribosome into 60S and 40S subunits, enabling subsequent dissociation of deacylated tRNA and mRNA from the 40S subunit and ensuring efficient recycling of ribosomes and mRNAs.

Although translation termination is generally efficient, the three stop codons exhibit varied efficiencies across individual mRNAs ^5^. Notably, UGA – the least efficient stop codon – is also the most frequently used in the human transcriptome. When a stop codon is decoded by a near-cognate tRNA, stop codon readthrough (SCR) occurs, generating protein products with C-terminal extensions ^6^. Extensive work has demonstrated that the nucleotide context surrounding the stop codon strongly influences termination fidelity ^7,8^. For example, a cytosine at the +4 position enhances SCR, consistent with structural evidence that eRF1 accommodates four nucleotides within the A site ^9^. In addition, RNA secondary structures downstream of stop codons can stimulate readthrough, as exemplified by the *kelch* mRNA in *Drosophila* ^10^. Such structures have been proposed to induce ribosomes pausing at stop codons, thereby biasing competition toward aminoacyl-tRNA decoding.

Beyond SCR, ribosome pausing at stop codons can also trigger frameshifting, as observed during translation of *OAZ* ^11^. Unlike SCR, which preserves the original reading frame, termination-coupled frameshifting produces protein products with heterogeneous C-terminal extensions. Promiscuous translation in 3’UTR has also been documented in cells lacking ABCE1 ^12^, and terminating ribosomes can undergo undergo reinitiation through bi-directional migration in a sequence-dependent manner ^13^. Together, these diverse outcomes underscore the dynamic nature of ribosome behavior o at stop codons and suggest that termination fidelity is tightly coupled to ribosome dynamics. However, very little is known whether individual stop codons exhibit distinct termination kinetics, and if so, what is the underlying mechanism.

Ribosome profiling provides a powerful approach to globally interrogate translation termination across all stop codons in their native sequence contexts ^14^. Application of this technique to diverse cell types has revealed that translational readthrough is more widespread in cellular mRNAs than previously appreciated ^15^. Intriguingly, readthrough displays marked tissue specificity, occurring at elevated levels in the central nervous system ^16,17^, whereas tissues such as testis exhibit minimal readthrough ^10^. These observations suggest that termination fidelity is governed not only by *cis*-acting mRNA elements but also by *trans*-acting factors that modulate the behavior of terminating ribosomes. The mechanistic basis of such cell type-specific regulation, however, remains poorly understood.

Ribosome stalling at premature termination codons has long been associated with nonsense-mediated mRNA decay ^18^. However, this model has been challenged by recent studies reporting comparable ribosome occupancy at premature and normal termination codons ^19^. Given that nonsense mutations account for >10% of inherited human diseases, including cystic fibrosis and muscular dystrophy ^20^, there is strong therapeutic interest in selectively promoting readthrough at premature stop codons while preserving accurate termination at normal stop codons. Achieving this goal requires a deeper understanding of ribosome dynamics and regulatory mechanisms operating at individual termination sites.

Here, by profiling terminating ribosomes in mammalian cells and mouse tissues, we uncover a broad spectrum of termination dynamics across the transcriptome. We show that the dwell time of terminating ribosomes is shaped not only by local sequence context but also by ribosome composition. Unexpectedly, ribosome pausing at stop codons functions as a protective translational checkpoint that restrains ribosome sliding and suppresses aberrant translation into the 3’UTR. Mechanistically, we identify the underlying mechanism by uncovering sequence-dependent interactions between mRNA and rRNA that underlie termination pausing, revealing an unanticipated layer of translational control with important physiological implications.

## RESULTS

### High-resolution Ribo-seq reveals dynamic features of terminating ribosomes

Ribosome profiling provides a global snapshot of translation by sequencing ribosome-protected mRNA fragments (RPFs) ^21^. Relative to the coding sequence (CDS), ribosome footprints accumulate at both start and stop codons. We recently developed Ezra-seq, a high-resolution ribosome profiling method with exceptional 5’ end precision ^22^. Owing to its superior 3-nt periodicity, Ezra-seq enables sensitive detection of start codon-associated ribosome frameshifting ^22^. We reasoned that Ezra-seq would also allow detailed characterization of terminating ribosomes. Indeed, when mRNAs are aligned to their annotated stop codons, we observed sharply defined boundaries of termination-associated footprints (Figure 1A, right panel). Unlike initiating ribosomes, which position the AUG codon in the P site, terminating ribosomes place the stop codon in the A site, as indicated by a prominent 5’ end peak at –15 nt relative to the stop codon. Consistent with previous reports ^14^, terminating ribosomes protected a longer stretch of mRNA (Figure 1A, bottom panel). Compared with elongating ribosomes, terminating ribosomes displayed two distinct footprint populations: long (30 – 31 nt) and short (20 – 23 nt) reads, a feature not observed for initiating ribosomes (Figure 1A, bottom panel). These shorter footprints likely represent terminating ribosomes with an unoccupied A-site.

**Fig. 1.**
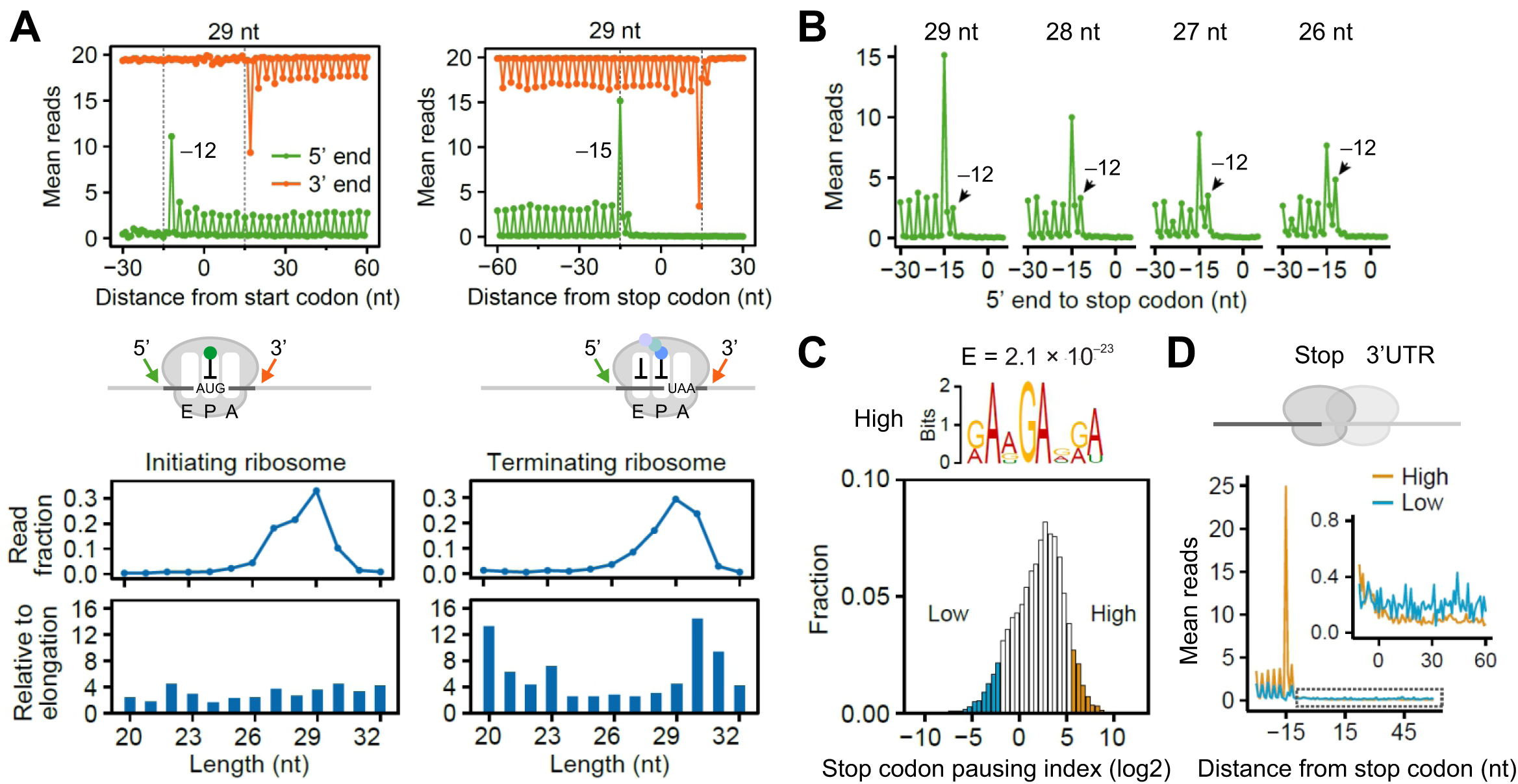
Characterizing terminating ribosomes using Ribo-seq. **(A)** Aggregation plots of mean Ribo-seq reads around the start (left) and stop (right) codons. Both 5’ end (green) and 3’ end (orange) of footprint reads (29 nt) were used for mapping. For bottom panels, the line plots show the distribution of reads with different lengths, whereas the bar plots show the read distribution of initiating (left) and terminating (right) ribosomes relative to elongation ribosomes. **(B)** Aggregation plots of mean Ribo-seq reads around the stop codon. The reads were stratified by the length followed by mapping using the 5’ end of footprint reads. **(C)** A histogram plot of ribosome pausing index at individual stop codon. The top shows the enriched sequence motif of mRNAs with strong ribosome pausing at stop codons. **(D)** Aggregation plots of mean Ribo-seq reads around the stop codon. **“**High” and “Low” refer to mRNAs with differential pausing indexes shown in (C). The 3’UTR read density was highlighted in the insert.

Closer inspection of stop codon-associated footprints revealed an additional 5’ end peak at –12 nt, which became more prominent among shorter reads (Figure 1B). Previous toe-printing assays have shown that eRF1 induces a forward movement of terminating ribosomes, shifting the leading edge from +13 nt to +15 nt ^23^. Moreover, single-molecule analyses have identified distinct pre- and post-termination phases catalyzed by eRF1 ^24^. Together, these observations suggest that the two 5’ end peaks correspond to pre- and post-terminating ribosome states, with the latter likely adopting a rotated conformation. However, we cannot exclude the possibility that a subset of these ribosomes transiently positions the stop codon in the P-site prior to ribosome disassembly.

To probe the kinetics of termination pausing, we performed cycloheximide (CHX) chase experiments in HEK293 cells. CHX stalls elongating ribosomes on mRNAs but does not arrest terminating ribosomes ^25^. Whereas a 5 min CHX treatment substantially reduced footprint density at stop codons, a 30 min pre-treatment nearly abolished the stop codon-associated peaks (Figure S1A and S1B). These results indicate that terminating ribosomes undergo transient pausing rather than stable stalling at the stop codon. In line with this notion, we observed little accumulation of upstream ribosomes, suggesting that termination pausing rarely triggers ribosome collisions.

Despite the overall presence of termination pausing, ribosome density at individual stop codons varied over several orders of magnitude. To quantify this variability, we calculated a stop codon pausing index for each mRNA by normalizing read density at the stop codon to average CDS occupancy, thereby accounting for differences in mRNA abundance and translation efficiency (Figure 1C). To identify sequence elements associated with termination pausing, we compared mRNAs with high and low pausing indices. Although no significant motifs were detected downstream of stop codons, a GA-rich motif upstream of the stop codon was strongly enriched among mRNAs exhibiting robust termination pausing (*E* = 2.1 × 10^−23^, Figure 1C). These findings indicate that certain coding sequences upstream of the stop codon play a critical role in modulating termination dynamics.

For mRNAs lacking termination pausing, ribosomes may either dissociate i rapidly from the mRNA or undergo stop codon readthrough. These scenarios are expected to produce distinct ribosome occupancy patterns in 3’UTR. Strikingly, we observed higher 3’UTR ribosome density on mRNAs lacking termination pausing (Figure 1D). This effect could not be explained by downstream sequence bias, as the identity of the +4 nt had minimal impact on 3’UTR translation (Figure S1C). Moreover, 3’UTR-associated footprints exhibited poor 3nt periodicity, in sharp contrast to the well-phased CDS reads (Figure S1D). These observations suggest that canonical stop codon readthrough alone cannot account for the elevated 3’UTR ribosome density observed in the absence of termination pausing.

### Profiling of terminating ribosomes by eRF1-seq

To directly interrogate the dynamics of translation termination, we developed a terminating ribosome profiling strategy by isolating ribosomes associated with eRF1 (Figure 2A). HEK293 cells were first crosslinked by formaldehyde, followed by cell lysis and RNase I digestion. Monosomes were isolated by sucrose gradient centrifugation, and eRF1-bound ribosomes were subsequently enriched by immunoprecipitation, as confirmed by immunoblotting (Figure S1E). In the absence of crosslinking, ribosomal proteins were minimally recovered by the eRF1 antibody, consistent with the transient nature of eRF1-ribosome interactions. Deep sequencing of ribosome-protected fragments from eRF1-bound ribosomes (eRF1-seq) revealed a pronounced accumulation of reads precisely at the annotated stop codons (Figure 2A, right panel). Remarkably, eRF1-seq retained single-nucleotide resolution, enabling unambiguous identification of termination sites on endogenous mRNAs (Figure 2B). Similar to conventional Ribo-seq, eRF1-seq also observed the forward-shifted footprints characteristic of post-termination ribosomes (Figure 2C). With excellent reproducibility between biological replicates (Figure S1F), eRF1-seq robustly and selectively profiles terminating ribosomes.

**Fig. 2.**
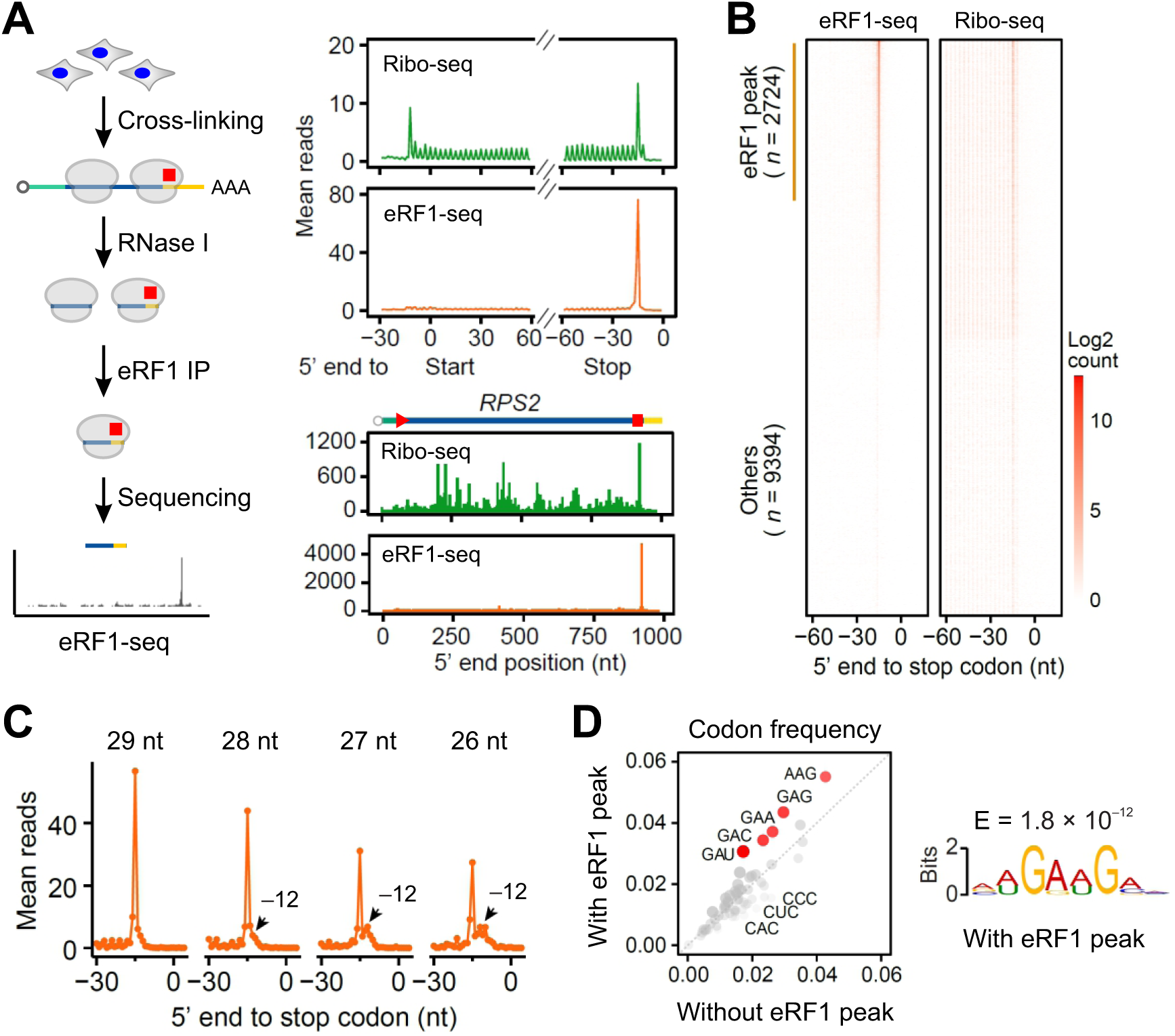
Characterizing terminating ribosomes using eRF1-seq. **(A)** The left panel shows the schematic of eRF1-seq procedures. The right top panel shows the aggregation plots of Ribo-seq (green) and eRF1-seq (orange). The right bottom panel shows a representative mRNA (*RPS2*) with reads obtained from Ribo-seq and eRF1-seq. The 5’ end of reads were used for mapping. **(B)** Heatmaps of individual mRNAs with reads obtained from Ribo-seq (right) and eRF1-seq (left). **(C)** Aggregation plots of mean eRF1-seq reads around the stop codon. The reads were stratified by the length followed by mapping using the 5’ end of footprint reads. **(D)** Comparison of codon frequencies upstream of stop codons between mRNAs with and without eRF1 peaks. The right panel shows the enriched sequence motif for mRNAs with eRF1 peaks at stop codons.

Consistent with our Ribo-seq analyses, eRF1-seq revealed substantial heterogeneity in termination pausing, as not all mRNAs displayed prominent eRF1 peaks at their annotated stop codons (Figure 2B). To identify determinants of this variability, we compared mRNAs with and without eRF1 peaks. Neither 3’UTR length nor downstream sequence features correlated with eRF1 peak height (Figure S1G). In contrast, sequences upstream of the stop codon were significantly enriched for a GA-rich motif among mRNAs exhibiting strong eRF1 peaks (Figure 2D). This observation mirrors the motif identified by Ribo-seq analysis (Figure 1C), reinforcing the conclusion that termination pausing is an intrinsic, sequence-encoded property of individual mRNAs. Notably, all three stop codons exhibited similar pausing features and shared upstream sequence motifs (Figure S1G and S1I).

### eRF1-seq reveals alternative termination sites

Despite the prominent peak at annotated stop codons, individual eRF1 peaks were broadly distributed across transcripts, spanning the 5’UTR, CDS, and 3’UTR (Figure 3A). The presence of eRF1 peaks in the 5’UTR is not unexpected, as ∼50% of human mRNAs harbor upstream open reading frames (uORFs) ^26^. Indeed, ∼30% of eRF1 located in the 5’UTR coincided with termination sites of uORFs previously identified by Ribo-seq (Figure S2A). By revealing additional, previously unannotated termination sites in the 5’UTR, eRF1-seq substantially expands the catalog of uORFs. Nearly all A-site codons underlying 5’UTR eRF1 peaks corresponded to canonical stop codons (Figure 3B), and Ribo-seq revealed a reduction in read density downstream of those sites (Figure 3B, right panel), consistent with *bona fide* translation termination. In addition to the 5’UTR, we identified a total of 807 eRF1 peaks in the 3’UTR that shared similar features (Figure S2B). Compared with conventional Ribo-seq, which typically yields sparse 3’UTR reads, eRF1-seq provided an improved signal-to-noise ratio for detecting translation events in the 3’UTR.

**Fig. 3.**
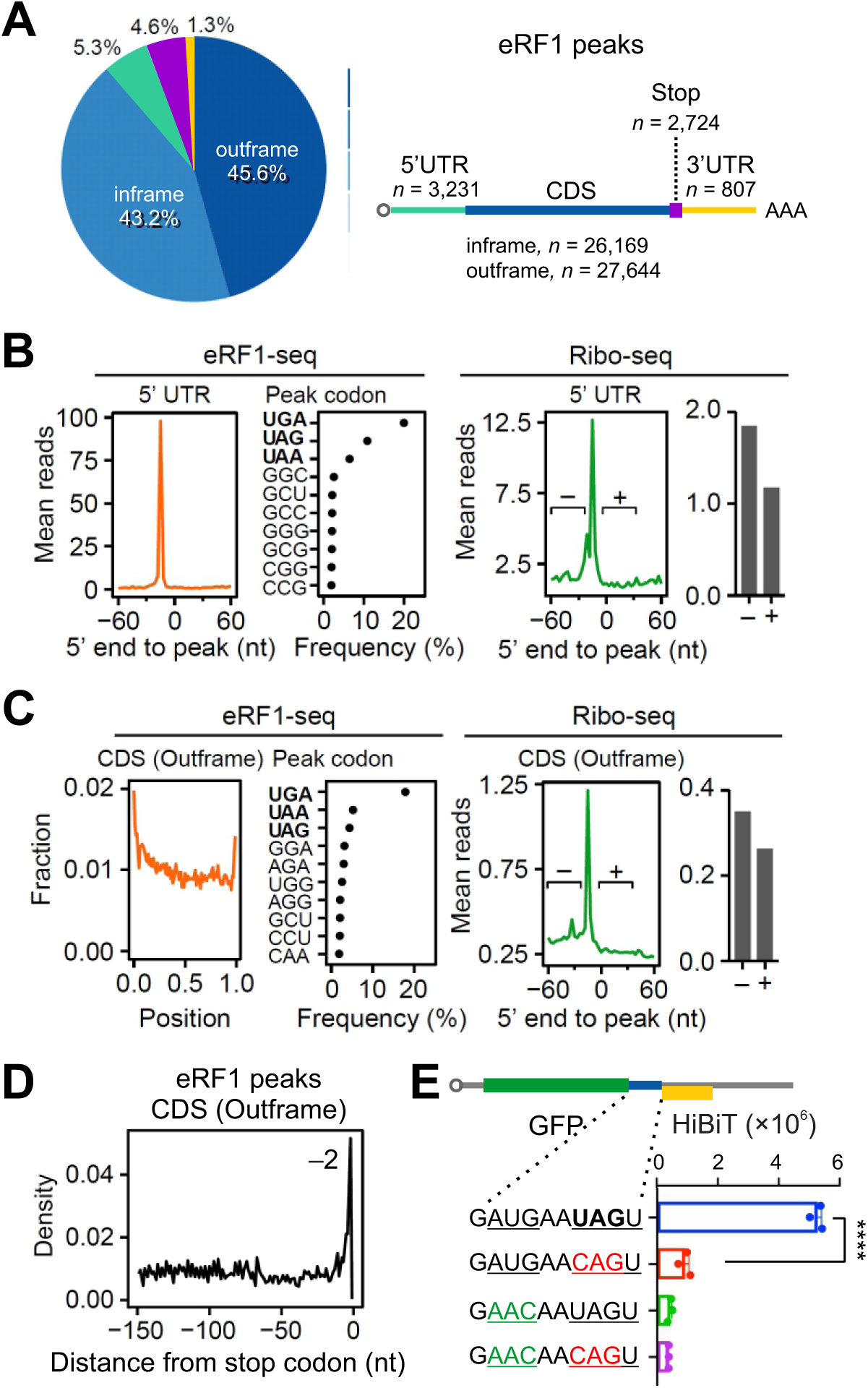
eRF1-seq reveals prevailing termination sites. **(A)** A pie chart shows fractions of eRF1 peaks mapped to different mRNA regions. “Inframe” and “Outframe” refer to the positions of eRF1 peaks within CDS relative to the annotated start codons. **(B)** The left panel shows mean eRF1-seq reads around the position of eRF1 peaks within 5’ UTR. The dot plot shows the frequency of A-site codons at eRF1 peaks. The right panel shows mean Ribo-seq reads around the position of eRF1 peaks within 5’ UTR. The bar graph shows the mean ribosome densities before (-) and after (+) the eRF1 peaks. **(C)** The left panel shows mean eRF1-seq reads around the position of out-of-frame eRF1 peaks within CDS. The dot plot shows the frequency of A-site codons at eRF1 peaks. The right panel shows mean Ribo-seq reads around the position of out-of-frame eRF1 peaks within CDS. The bar graph shows the mean ribosome densities before (-) and after (+) the eRF1 peaks. **(D)** The distribution of eRF1 peaks before the annotated stop codon. Only the out-of-frame eRF1 peaks were used for plotting. **(E)** A bar graph shows the HiBiT-based reporter assays in HEK293-K^b^ cells. HiBiT signals were measured from cells transfected with mRNA reporters bearing different sequences between GFP and HiBiT. Error bars, mean ± s.e.m. *n* = 3 biological replicates. \*\*\*\**P* ≤ 0.0001 by unpaired two-tailed *t*-test.

Even under stringent peak-calling criteria, eRF1-seq detected a substantial number of peaks within CDS. Many of these CDS-associated eRF1 peaks positioned their A sites at in-frame sense codons (Figure S2C). Notably, the majority of those codons were “stop-like” in that they corresponded to frameshifted versions of canonical stop codons. For example, CUG represents a –1 frameshifted UGA, whereas GAG, GAA, and AAG represent +1 frameshifted UGA or UAA. Despite robust eRF1 occupancy, those sense codons did not trigger translation termination, as Ribo-seq revealed no decrease in downstream read density (Figure S2C). These observations suggest that eRF1 can transiently compete with cognate A-site tRNAs during elongation, resulting in false termination attempts.

In addition to in-frame sense codons, a large fraction of CDS-associated eRF1 peaks corresponded to out-of-frame canonical stop codons, particularly UGA (Figure 3C). Supporting authentic termination at these sites, Ribo-seq revealed reduced ribosome occupancy downstream (Figure 3C, right panel). Interestingly, most out-of-frame termination events occurred near the beginning of CDS. Beyond overlapping uORFs, this pattern further supports the occurrence of start codon-associated ribosome frameshifting that we reported recently ^22^. Out-of-frame eRF1 peaks were also enriched near the 3’ end of CDS (Figure 3C, 3D), consistent with migration of terminating ribosomes at stop codons to search for upstream start codons (Figure 3D). To directly validate stop codon-associated reinitiation, we inserted a 9 nt sequence derived from *CASQ2*, which contains an out-of-frame AUG codon upstream of a UAG stop codon, between GFP and HiBiT (Figure 3E). Consistent with previous reports ^27^, mutation of the UAG stop codon abolished reinitiation and eliminated out-of-frame HiBiT expression (Figure 3E), confirming that reinitiation is dependent on termination at the stop codon.

### Sequence determinants of termination pausing

Given the wide range of termination pausing observed at individual stop codons, we next thought to identify the sequence determinants of termination pausing using a massively paralleled reporter assay (MPRA). In contrast to analysis of endogenous genes, whose sequences are constrained by evolutionary selection, MPRA leverages fully randomized sequences to enable unbiased identification of sequence elements governing translation behavior. We previously employed a uORF-based MPRA to assess start codon usage with randomized sequence contexts ^28^. In this system, mRNA variants with efficient uORF translation are retained in monosome fraction, whereas efficient translation of downstream GFP relocates the mRNA to polysome. Thus, the monosome/polysome ratio serves as a quantitative readout of uORF translation efficiency.

To interrogate stop codon usage, we modified the uORF reporter by replacing the uORF stop codon with a 9-nt randomized sequence (Figure 4A). Introduction of an in-frame stop codon within the insert would terminate uORF translation and restrict the mRNA to the monosome fraction. To eliminate transcriptional variation associated with plasmid-based expression, we synthesized the reporter library by *in vitro* transcription. Following transfection of the mRNA pool into HEK293 cells, monosome and polysome fractions were separated by sucrose gradient centrifugation and analyzed by deep sequencing. Among mRNAs recovered from the monosome fraction, all three canonical stop codons (UGA, UAG, UAA) were strongly enriched (Figure 4A). Importantly, these stop codons were preferentially positioned in-frame within the randomized insert (Figure S3A and S3B). Codons enriched in alternative reading frames were also informative; for example, codons enriched in frame 2 predominantly belong to NUA and NUG, consistent with frameshifted presentations of in-frame stop codons (Figure S3B, bottom panel). Together, these results establish MPRA as a robust platform for dissecting stop codon-associated sequence features.

**Fig. 4.**
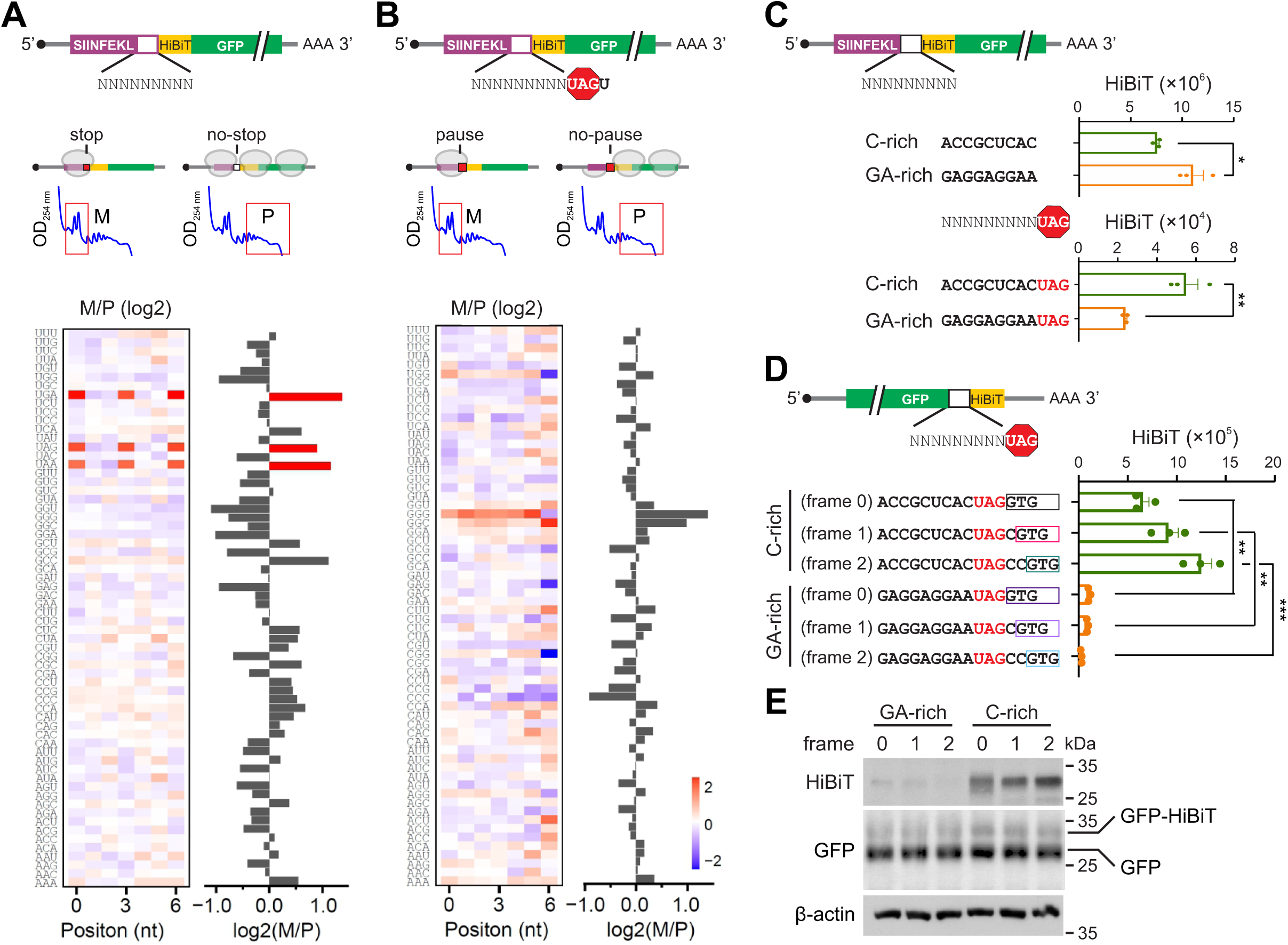
Termination pausing is influenced by sequence contexts. **(A)** The up panel shows the schematic of a massively parallel reporter assay, where the stop codon of uORF was replaced with a random 9-nt sequences. The uORF translation was monitored by the number of associated ribosomes separated by sucrose gradient (M, monosome; P, polysome). The heatmap shows the ratio of monosome fraction over polysome fraction (M/P) when different codons were placed at individual positions within the 9-nt random sequence (left). The stop codons UGA, UAG and UAA are highlighted. The bar graph (right) shows the mean M/P ratio averaged across all positions. **(B)** Similar as (A) except that the random 9-nt sequences were placed before the stop codon UAG. **(C)** Bar graphs show the HiBiT signals in HEK293-K^b^ cells transfected with mRNA reporters bearing C-rich or GA-rich sequences between the uORF and HiBiT-GFP. The up panel shows the control without the stop codon UAG. Error bars, mean ± s.e.m. *n* = 3 biological replicates. \**P* ≤ 0.05; \*\**P* ≤ 0.01 by unpaired two-tailed *t*-test. **(D)** A bar graph shows the HiBiT signals in HEK293-K^b^ cells transfected with mRNA reporters bearing C-rich or GA-rich sequence before the GFP stop codon UAG. The downstream HiBiT sequence was inserted into different reading frames. Error bars, mean ± s.e.m. *n* = 3 biological replicates. \*\**P* ≤ 0.01; \*\*\**P* ≤ 0.001 by unpaired two-tailed *t*-test. **(E)** Representative Western blots of GFP and HiBiT in HEK293-K^b^ cells transfected with C-rich or GA-rich reporters as described in (D).

Ribosomal pausing at uORF stop codons is expected to retain mRNAs in the monosome fraction by suppressing both leaky scanning and stop codon readthrough. We therefore leveraged the monosome/polysome ratio as a proxy for termination pausing. We first examined the impact of downstream sequences by placing a 9-nt randomized sequence immediately after a fixed UAG stop codon (Figure S3C). Despite prior reports that a cytosine (C) at the +4 nt position enhances readthrough ^7^, mRNA variants containing C-rich downstream sequence were depleted from both monosome and polysome fractions. This depletion likely reflects accelerated mRNA turnover caused by translation into 3’UTR ^29,30^. Consistent with our Ribo-seq and eRF1-seq analyses, no specific downstream sequences were enriched in the monosome fraction (Figure S3D), indicating that sequences following the stop codon exert minimal influence on termination pausing.

We next assessed whether upstream sequences modulate stop codon fidelity by inserting a 9-nt randomized sequence immediately before the UAG stop codon (Figure 4B). Strikingly, G-rich sequences were strongly enriched in the monosome fraction (Figure 4B and S3E). Notably, this enrichment lacked reading frame information, indicating that nucleotide composition rather than codon identity governs ribosome pausing at stop codons. In contrast, C-rich sequences were selectively depleted from the monosome fraction (Figure 4B), suggesting that the C-rich upstream contexts promote sequence stop codon readthrough and downstream translation. These findings closely mirror the sequence dependencies revealed by Ribo-seq and eRF1-seq, reinforcing the conclusion that G-rich sequence intrinsically favor termination pausing.

### Stop codon-associated random translation

It was not immediately clear whether translation in the 3’UTR arose from stop codon readthrough or stop codon-associated reinitiation. To distinguish these possibilities, we investigated the nature of downstream translation triggered by upstream C-rich sequences using individual mRNA reporters. Specifically, we placed a C-rich sequence immediately upstream of the uORF stop codon UAG (Figure 4C). Because many C-rich triplets encode proline, an imino acid known to affect ribosome dynamics, we deliberately selected non-proline C-rich codons to avoid confounding effects. As a control, we used a GA-rich sequence identified from eRF1-seq as being associated with strong termination pausing. To sensitively detect translation downstream of the stop codon, we inserted a HiBiT coding sequence immediately after UAG and HiBiT signals can be detected with superior sensitivity ^28^. The canonical start codon of HiBiT was omitted to exclude leaky scanning-mediated translation. In the absence of the stop codon, the GA-rich sequence produced higher HiBiT signals than the C-rich sequence, likely reflecting difference in the amino acids encoded by the inserts. Strikingly, in the presence of the UAG stop codon, the C-rich sequence nearly doubled HiBiT signals compared with the GA-rich sequence (Figure 4C, bottom panel). This result is consistent with the MPRA data, indicating that C-rich coding sequences preceding the stop codon not only attenuate termination pausing but also actively promote downstream translation. Importantly, this phenomenon was not restricted to uORFs, as similar results were obtained when the uORF was replaced by a GFP coding sequence (Figure S3F).

Canonical stop codon readthrough is expected to proceed in-frame and generate a fusion protein with a C-terminal extension. To test this model, we placed the HiBiT sequence in different reading frames relative to the stop codon. Unexpectedly, robust HiBiT signals were detected from reporters in both frame 1 and frame 2 (Figure 4D). In contrast, the GA-rich sequence preceding the stop codon strongly suppressed downstream HiBiT translation. This frame-independent effect was reproducible across multiple sequence variants (Figure S3G). To directly examine the translational products, we conducted immunoblotting of whole cell lysates using HiBiT antibodies (Figure 4E). GFP-fusion proteins (∼ 30 kDa) were readily detected in C-rich reporters irrespective of the HiBiT reading frames, ruling out reinitiation as the underlying mechanisms. Notably, levels of non-fusion GFP were comparable across reporters, indicating that the differential HiBiT signals were not attributable to altered translation initiation or differences in the amino acids encoded by the upstream inserts. Together, these results demonstrate that C-rich coding sequence promote ribosome sliding at the stop codon, enabling translation into the 3’UTR in all three reading frames.

We next examined the positional dependence of the C-rich effect by introducing C triplets at varying distance upstream of the stop codon. HiBiT reporter assays revealed that a C triplet positioned immediately adjacent to the stop codon was the least effective at promoting 3’UTR translation (Figure S3H). This finding suggests that the sequence determinant governing ribosome sliding resides upstream of the E site, rather than directly at the termination codon itself.

### ABCE1 regulates terminating ribosomes independent of the sequence context

ABCE1 (Rli1 in yeast) is a conserved ABC-type protein that plays a crucial role in translation termination and ribosome recycling ^3^. Previous studies have shown that depletion of ABCE1 leads to ribosome stalling at stop codons and increased ribosome occupancy within 3’UTRs ^12^. To investigate whether ABCE1 exhibits sequence preference towards terminating ribosomes, we knocked down ABCE1 in HEK293 cells using shRNA (Figure S4A). As expected, ABCE1 silencing impaired cell proliferation and reduced global protein synthesis (Figure S4B). We next performed ribosome profiling in ABCE1 KD and control cells using Ezra-seq. After normalizing ribosome occupancy across CDS, loss of ABCE1 resulted in a modest but reproducible increase in ribosome density at stop codons (Figure S4C). Notably, this increase was observed at all three stop codons, an indication of a global effect. A closer inspection of footprint distribution revealed that ABCE1 depletion selectively increased ribosome density at the –15 nt position, whereas the forward-shifted –12 nt peak was largely unaffected (Figure S4D). These results suggest that ABCE1 primarily facilitates late-stage termination or ribosome splitting, and its absence delays pre-termination progression. A prior study reported that ABCE1 knockdown in HeLa cells promotes translation into 3’UTR in all three reading frames ^12^. We were unable to robustly detect this phenomenon by Ribo-seq, likely due to the limited sequencing depth and/or incomplete ABCE1 knockdown in HEK293 cells (Figure S4A). To overcome these limitations, we employed a sensitive HiBiT-based 3’UTR reporter assay, which showed increased 3’UTR translation irrespective of the sequence context upstream of the stop codon (Figure S4E). Therefore, the ribosome recycling factor ABCE1 plays a generic role in translation termination independent of mRNA sequence context.

### The 3’ end of 18S rRNA influences termination pausing

For a terminating ribosome, the mRNA sequence preceding the stop codon is positioned upstream of the E-site. Given the narrow dimensions of the mRNA channel, it is unlikely that external protein factors such as ABCE1 directly regulate ribosome behavior in a sequence-specific manner. Instead, prior crosslinking studies have shown that mRNA near the exit site interacts with the 3’ terminus of 18S rRNA ^31^. Such proximity is lso evident in recent cryoEM structures of mammalian initiating ribosomes ^32^, although stable base pairing was not explicitly resolved (Figure 5A). A similar juxtaposition between mRNA and the 3’ end of 18S rRNA has been observed in elongating ribosomes during translocation ^33^. Structural modeling based on available cryo-EM data suggests that, from early to late translocation intermediates (POST-1 to POST-3), the distance between the mRNA segment (– 9 to – 3) and the 3’ end of 18S rRNA progressively decreases (Figure 5B). These observations support a model in which mRNA undergoes post-decoding scanning by the 3’ terminus of 18S rRNA as it exits the ribosome. Notably, the 3’ end of 18S rRNA harbors a highly conserved U-rich sequence (GAUCAUUA). We propose that GA-rich mRNA sequences can engage in transient U:A and U:G base pairing with 18S rRNA near the exit site (Figure 5A and S5A), thereby slowing mRNA passage and promoting termination pausing. In contrast, C-rich sequences would evades this rRNA-mediaed checkpoint, resulting in faster mRNA transit and reduced termination pausing.

**Fig. 5.**
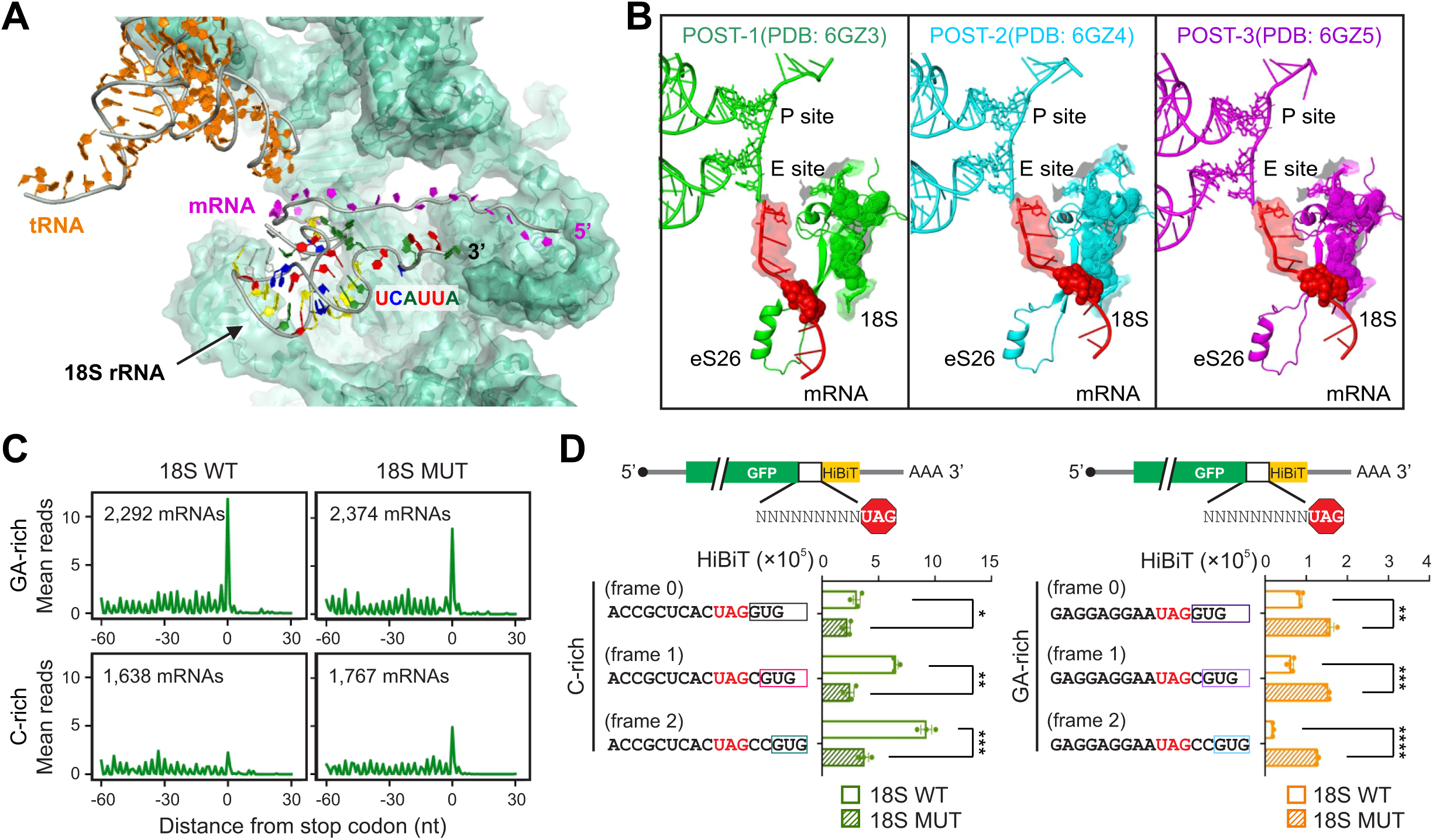
The 3’ end of 18S rRNA influences the dynamic of terminating ribosome. **(A)** A cryoEM structures of mammalian initiating ribosomes (PDB: 6ZMW) showing the proximity of the 3’ terminus of 18S rRNA and mRNA. **(B)** Superposition of different conformations sampled by normal mode analysis (NMA) of the region mentioned in (A). The analysis was carried out with the mRNA segment from the -10 nt to the +5 nt related to the A-site. The superposition is outlined only for the Rps26 protein (green surface), the mRNA segment (blue surface) and the last 10 nucleotides at the 3’ rRNA extremity (dark red surface), with the remaining proteins and rRNA segments at the site described only by the average structure (silver transparent surface and cartoon). The tRNA anticodon loop at the P ribosomal site is shown in dark cyan. **(C)** HEK293 cells were transfected with plasmids encoding 18S rRNA WT or mutant followed by Ribo-seq. Aggregation plots show the mean reads around stop codons of mRNAs with the GA sequence motif or C-rich sequence element. **(D)** HEK293 cells were transfected with plasmids encoding 18S rRNA WT or mutant followed by transfection with mRNA reporters shown in (4D). Bar graphs show the HiBiT signals in transfected cells with HiBiT at different reading frames. Error bars, mean ± s.e.m. *n* = 3 biological replicates. \**P* ≤ 0.05; \*\**P* ≤ 0.01; \*\*\**P* ≤ 0.001; \*\*\*\**P* ≤ 0.0001 by unpaired two-tailed *t*-test.

To directly test whether putative mRNA: rRNA base pairing contributes to termination pausing, we sought to perturb the 3’ end sequence of 18S rRNA. Because the presence of hundreds of rDNA copies precludes genome editing approaches such as CRISPR/Cas9, we instead used a previously described 18S rRNA expression system that enables incorporation of exogenously expressed 18S rRNA into ∼15% of 40S subunits in transfected cells ^34,35^. We engineered an 18S rRNA mutant by substituting the last two U residues with G (GAUCAGGA), a change predicted to weaken interactions with GA-rich mRNA while favoring pairing with C-rich sequence. Overexpression of the mutant 18S in HEK293 cells resulted in polysome profiles comparable to those observed with wild-type 18S rRNA (Figure S5B), and ribosome profiling by Ezra-seq revealed similar ribosome occupancy across coding regions (Figure S5C). However, when mRNAs were stratified by the sequence motif upstream of the stop codon, expression of the mutant 18S reduced the differential termination pausing between GA-rich and C-rich sequences (Figure 5C). Specifically, mRNAs containing GA-rich motifs exhibited reduced stop codon peaks, whereas those bearing C-rich motifs showed increased ribosome accumulation at stop codons. The reciprocal shift in termination pausing in response to mutation of the 18S rRNA 3’ end provides strong evidence that sequence-specific termination pausing is mediated by interaction between mRNA and 18S rRNA.

We further validated this model using HiBiT-based reporters to assess 3’UTR translation in cells expressing WT or mutant 18S rRNA. Expression of the mutant 18S rRNA attenuated 3’UTR translation from reporters containing C-rich upstream sequences (Figure 5D). Conversely, reporters bearing GA-rich sequences exhibited increased HiBiT signals, indicative of enhanced 3’UTR translation. Together with the swapped termination pausing profiles observed by ribosome profiling, these results establish a crucial role for the 3’ end of 18S rRNA in enforcing termination fidelity. Notably, the 3’ terminal sequence of 18S rRNA is highly conserved across eukaryotes (Figure S5D). Annotated stop codons in the human genome are preferentially preceded by GA-rich sequences (Figure S5E). In contrast, out-of-frame stop codons show a higher prevalence of C-rich upstream contexts. These patterns suggest an evolutionary advantage for termination pausing at annotated stop codons, likely serving to minimize stop codon slippage and aberrant readthrough while preserving proteome integrity.

### Tissue-specific termination pausing

Having established sequence-specific termination pausing in cultured cell lines, we next explored its physiological relevance across tissues. To this end, we conducted Ribo-seq on a diverse panel of mouse tissues, including liver, brain, kidney, heart, and testis (Figure 6A). These tissues displayed distinct polysome profiles and global ribosome occupancy across coding regions (Figure 6A and S6A), reflecting substantial differences in translational states. Strikingly, ribosome density at start and stop codons varied most prominently among tissues and did so in a reciprocal manner. Testis exhibited relatively modest initiation peaks but pronounced ribosome accumulation at stop codons (Figure 6A). In contrast, liver, heart, and brain showed robust initiation peaks with little or no detectable termination pausing. These differences were not attributable to tissue-specific gene expression, as the same reciprocal pattern persisted when analyses were restricted to genes commonly expressed or uniquely expressed in liver and testis (Figure S6B). Consistent with sequence-dependent termination pausing, mRNAs with prominent stop codon peaks in testis were enriched with GA-rich sequences upstream of the stop codon (Figure S6C). Notably, these same mRNAs exhibited minimal termination pausing in liver. Moreover, the attenuated stop codon peak in mouse liver has been independently reported ^36^, excluding the possibility of technical bias in Ezra-seq.

**Fig. 6.**
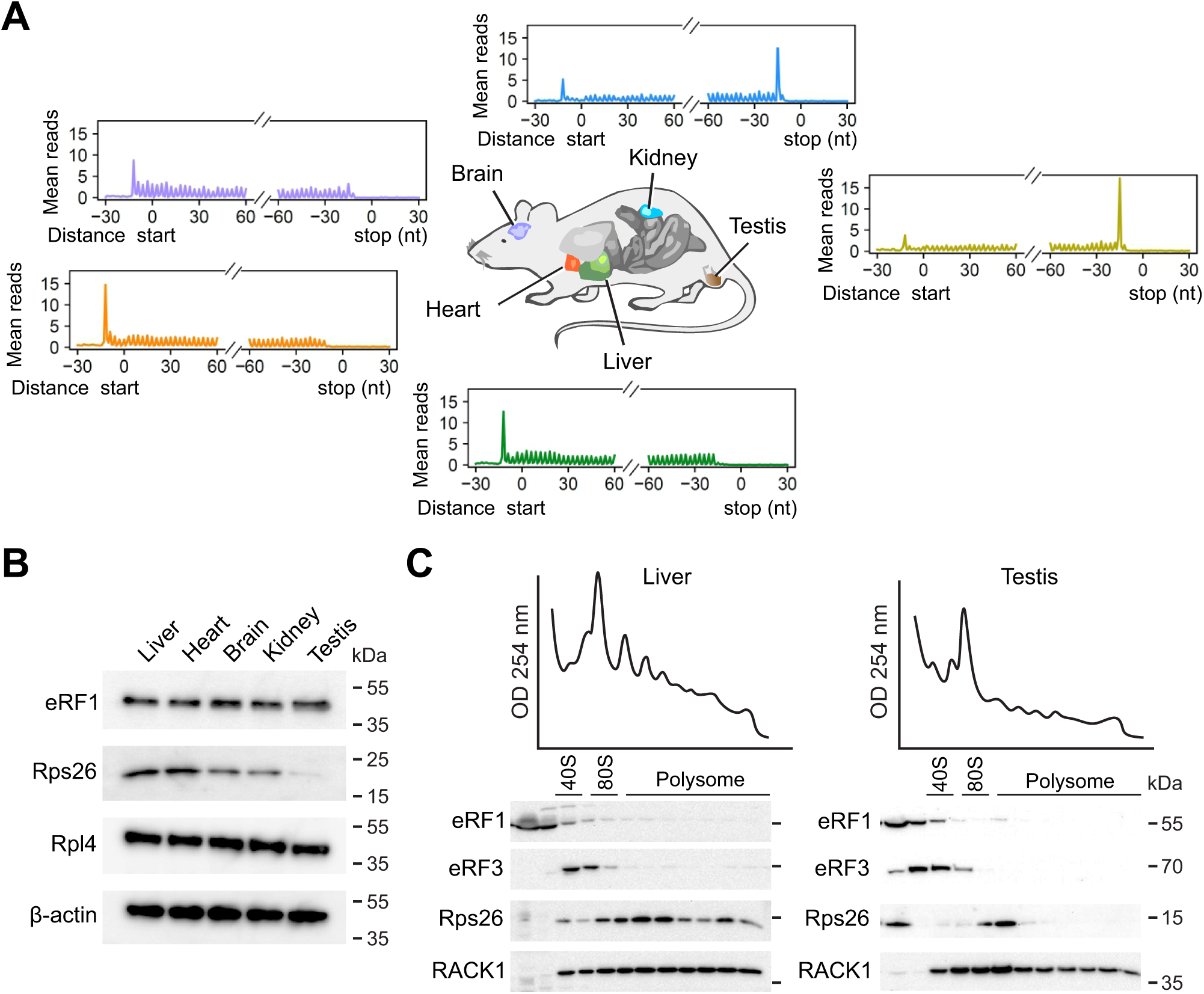
Differential termination pausing in mouse tissues. **(A)** Different mouse tissues were collected followed by Ribo-seq. Metagene analysis shows the distribution of mean ribosome reads across the transcriptome aligned to start and stop codons. **(B)** Representative Western blots of different mouse tissues using antibodies indicated. The experiment was independently repeated three times with similar results. **(C)** Mouse liver and testis were subjected to polysome profiling using sucrose gradient. Representative Western blots of ribosome fractions were conducted using antibodies indicated. The experiment was independently repeated three times with similar results.

The unexpected tissue-specific variation in termination pausing points to an additional regulatory layer beyond mRNA sequence context. One candidate is the eukaryote-specific ribosomal protein S26 (Rps26), which contacts mRNA upstream of the E-site ^31,37^. Recent cryo-EM structures revealed that both Rps26 (AA62 – 70) and 18S rRNA (nt1857 – 1863) jointly shape the mRNA path near the ribosomal exit site (Figure 5B). Because Rps26 lies between the mRNA and the 3’ end of 18S rRNA, its absence could enhance mRNA: rRNA interactions and thereby strengthen termination pausing. Supporting this notion, a recent study demonstrated that Rps26 can dissociate from fully assembled 80S ribosomes under stress conditions, generating ribosome heterogeneity ^38^. To assess whether Rps26 abundance varies across tissues, we compared ribosomal proteins in tissue homogenates. When normalized to β-actin, core ribosomal proteins such as Rpl4 were present at comparable levels across tissues, whereas Rps26 levels were markedly reduced in testis relative to other tissues examined (Figure 6B). Differential Rps26 expression between liver and testis is also evident in Human Protein Atlas (proteinatlas.org) ^39^. We further examined Rps26 distribution across polysome fractions in liver and testis. In liver, Rps26 and the control ribosomal protein RACK1exhibited similar polysome association. In contrast, testis showed pronounced depletion of Rps26 from polysomes (Figure 6C), accompanied by accumulation of Rps26 in ribosome-free fractions. Whether this reflects active dissociation of Rps26 from translating ribosomes in testis remains to be determined. These findings nevertheless suggest Rps26 stoichiometry contributes to tissue-specific termination pausing, likely by modulating access of the mRNA to the 3’ end of 18S rRNA.

### Rps26 modulates termination pausing

Based on available ribosome structures containing bound mRNA, Rps26 is positioned between the exiting mRNA and the 3’ end of 18S rRNA (Figure 7A). We hypothesize that loss of Rps26 would increase the accessibility of the mRNA to the 3’ terminus of 18S rRNA, thereby facilitating mRNA:rRNA interactions and promoting termination pausing. Consistent with this model, normal mode analysis (NMA) using anisotropic network models predicts that removal of Rps26 permits coordinated twisting and closer approximate between the mRNA segment (–3 to – 9) and the 3’ end of 18S rRNA (Figure 7B). This enhanced spatial parity would be further stabilized when the mRNA sequence is complementary to the rRNA. Given the U-rich nature of the 18S rRNA 3’ end, such interactions are predicted to be most favorable for GA-rich mRNA sequences.

**Fig. 7.**
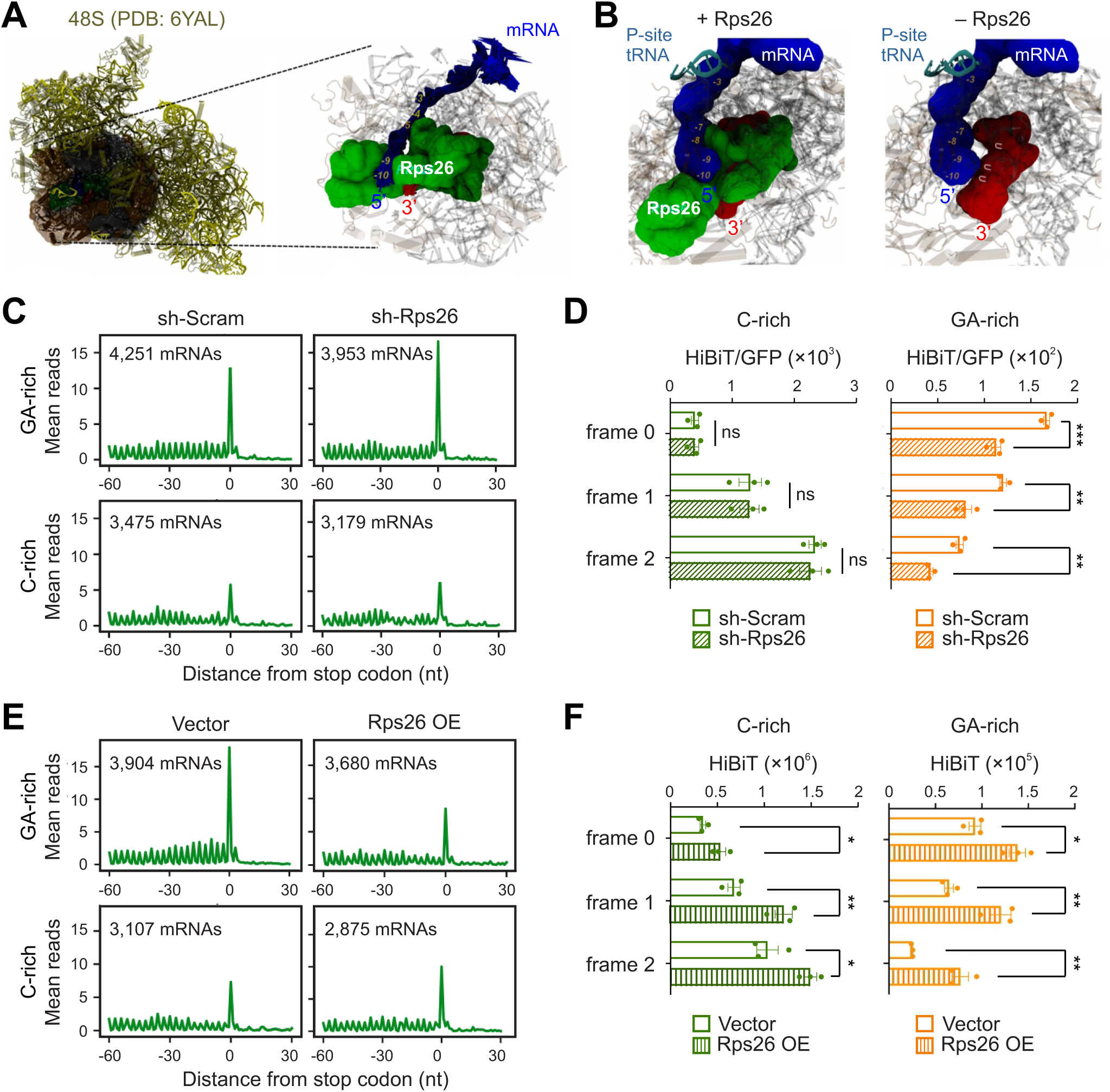
Rps26 modulates termination pausing. **(A)** A cryo-EM structure of the human 48S pre-initiation complex (PDB: 6YAL) depicted in cartoon and colored in yellow. The region at the 5’ exit of the mRNA tunnel centered in Rps26 and extended around 30 Å of this protein is highlighted in transparent surface and in dark colors. **(B)** Conformational sampling by normal mode analysis (NMA) of the region mentioned in (A). The analysis was carried out with the mRNA segment from the -10 nt related to the P-site to the +5 nt related to the A-site. The superposition is outlined only for the Rps26 protein (green surface), the mRNA segment (blue cartoon) and the last 10 nucleotides at the 3’ rRNA extremity (dark red cartoon), with the remaining proteins and rRNA segments at the site described only by the average structure (silver transparent surface and cartoon). The tRNA anticodon loop at the P ribosomal site is omitted on the image. **(C)** HEK293 cells with or without Rps26 knockdown were subjected to Ribo-seq. Aggregation plots show the mean reads around stop codons of mRNAs with the GA sequence motif or C-rich sequence element. **(D)** HEK293 cells with or without Rps26 knockdown were transfected with mRNA reporters shown in 4D. Bar graph shows the HiBiT signals at different reading frames upon normalization to upstream GFP levels Error bars, mean ± s.e.m. *n* = 3 biological replicates. ns, nonsignificant; \*\**P* ≤ 0.01; \*\*\**P* ≤ 0.001 by unpaired two-tailed *t*-test. **(E)** HEK293 cells with or without Rps26 overexpression were subjected to Ribo-seq. Aggregation plots show the mean reads around stop codons of mRNAs with the GA sequence motif or C-rich sequence element. **(F)** HEK293 cells with or without Rps26 overexpression were transfected with mRNA reporters shown in 4(D). Bar graph shows the HiBiT signals at different reading frames. Error bars, mean ± s.e.m. *n* = 3 biological replicates. \**P* ≤ 0.05; \*\**P* ≤ 0.01 by unpaired two-tailed *t*-test.

To test whether reduced Rps26 levels affect ribosome dynamics at stop codons, we depleted Rps26 in HEK293 cells using shRNA (Figure S7A). Rps26 silencing led to an increased 60S peak and reduced polysome formation (Figure S7B), consistent with its known role in 40S subunit maturation ^40^. We then conducted ribosome profiling in parallel with RNA-seq. Remarkably, Rps26 depletion resulted in reduced ribosome density at start codons but increased ribosome accumulation at stop codons (Figure S7C). This reciprocal change mirrors the tissue-specific differences observed between initiation and termination (Figure 6A). Importantly, the elevated termination pausing occurred predominantly at stop codons preceded with GA-rich sequences (Figure 7C). To assess the functional consequences of altered termination dynamics, we employed HiBiT-based 3’UTR reporter assay. Because Rps26 silencing globally reduced protein synthesis (Figure S7B), HiBiT signals were normalized to upstream GFP expression. Under these conditions, only reporters containing GA-rich sequences, but not C-rich sequences, exhibited reduced HiBiT translation in Rps26-depleted cells (Figure 7D). Therefore, Rps26 specifically modulates termination fidelity at GA-rich stop codons by constraining mRNA:rRNA interactions.

To further substantiate this conclusion, we overexpressed Rps26 in HEK293 cells (Figure S7D), which did not perturb polysome formation (Figure S7E). Ribosome profiling revealed that Rps26 overexpression lowered ribosome occupancy from initiation through termination (Figure S7F). When mRNAs are stratified by upstream sequence context, Rps26 overexpression ocaused a pronounced (>50%) reduction in ribosome density at stop codons preceded by GA-rich sequences (Figure 7E). Notably, ribosome density upstream of these stop codons were also diminished, suggesting that increased Rps26 levels broadly restrain mRNA:rRNA engagement during elongation. In contrast, mRNAs harboring the C-rich sequence upstream of the stop codon displayed relatively enhanced termination peaks. To directly link stop codon pausing with termination fidelity, we conducted HiBiT reporter assays t in cells with Rps26 overexpression. In line with reduced termination pausing, Rps26 overexpression increased 3’UTR translation (Figure 7F), confirming that diminished mRNA:rRNA interaction compromises termination fidelity. Collectively, these results establish Rps26 as a key modulator of sequence-specific termination pausing.

## DISCUSSION

Translation termination is intrinsically slower than elongation, and stop codons therefore frequently serve as ribosome pausing sites. Insufficient pausing at stop codons can lead to incomplete ribosome dissociation, allowing ribosomes to continue translating into the 3’UTR. Conversely, excessive pausing would delay ribosome recycling and increase the risk of ribosome collisions. Thus, ribosome dwell time at stop codons must be precisely calibrated to balance termination efficiency and fidelity. Since the advent of ribosome profiling, elevated ribosome density at both start and stop codons has been a consistent feature of metagene analyses. However, little has been known about the variability of termination pausing at individual stop codons, let alone the underlying mechanism. By combining high-resolution ribosome profiling ^22^ with a specialized eRF1-seq approach, we quantitatively defined stop codon pausing indices and uncovered a previously unappreciated role of mRNA sequence context in shaping the dynamics of terminating ribosomes. Unexpectedly, it is the sequence upstream of the stop codon – rather than the stop codon itself or downstream elements – that determine termination pausing. A GA-rich sequence element promotes ribosome pausing, a finding independently confirmed by massively paralleled reporter assay. In contrast, C-rich sequences upstream of stop codons abolish termination pausing. Importantly, loss of termination pausing does not simply result in canonical stop codon readthrough; instead, it triggers stop codon-associated random translation into the 3’ UTR, generating proteins with mixed C-terminal extensions. This phenomenon reveals a distinct and more complex mode of termination than classical readthrough.

A potential caveat in interpreting sequence-specific termination control is that different nucleotides sequences encode different amino acids, which could independently influence ribosome dynamics. This concern is particularly relevant for C-rich sequences, which often encode proline. However, multiple lines of evidence support the conclusion that nucleotide sequence – not amino acid identity – governs termination pausing. First, the positional effects of sequence elements do not conform to codon triplet boundaries. Second, although proline codons are decoded slowly, C-rich sequences reduce rather than enhance termination pausing. Third, C-rich sequence-promoted downstream translation persists even when proline codons are excluded. Fourth, fusion protein analyses reveal minimal effects of the encoded amino acids on translation outcomes. Fifth, unbiased MPRA analyses identify enrich sequence motifs across all reading frames. These findings collectively demonstrate that mRNA sequences, rather than the encoded peptides, control ribosome behavior at stop codons.

Because the mRNA segment upstream of the stop codon resides near the exit site of the ribosomal mRNA channel, our findings suggest that ribosome - mRNA communication extends beyond the decoding center. Such post-decoding interactions are well established during translation initiation, exemplified by the Shine-Dalgarno (SD) sequence that pairs with the 3’ end of 16S rRNA in prokaryotic cells ^41^. In eukaryotic cells, analogous base-pairing interactions involving the 3’ end of 18S rRNA has been documented in cap-independent translation and cap-dependent translation of histone H4 mRNA ^42–44^. We previously reported that the highly conserved 3’ terminus of 18S rRNA contributes to elongation pausing via base pairing with certain codons ^35^. Our current work extends this principle to translation termination, demonstrating that post-decoding mRNA:rRNA interactions near the exit channel modulate ribosome dwell time at stop codons.

Although ribosomal structures have been solved at near-atomic levels, relatively few contain well-resolved mRNA, owing to its intrinsic flexibility. Crosslinking studies have indicated proximity between mRNA and rRNA near the exit site ^31^, yet the functional relevance of this interaction remained unclear. Using targeted mutations of the 18S rRNA, we showed that U-rich sequence at its 3’ end is crucial for stabilizing mRNAs bearing GA-rich sequence motifs. The involvement of G:U base pairs is particularly notable, as these non-canonical interactions possess unique chemical properties distinct from A:U pairing ^45^. Mutation of the U-rich sequence to G reverses termination pausing preferences, shifting pausing from GA-rich to C-rich mRNAs. These results support a model in which mRNA:rRNA base pairing near the exit channel delays mRNA movement following decoding. For terminating ribosomes, the extended dwell time likely enhances eRF1 recruitment, peptide release, and efficient ribosome recycling.

While post-decoding mRNA:rRNA interactions explain sequence-specific termination pausing, they do not fully account for tissue-specific differences we observed. Termination pausing is particularly strong in testis, a tissue characterized by exceptional transcriptomic diversity arising from promiscuous transcription ^46^ and attenuated nonsense-mediated decay ^47^. Testis translation also displays unique features, including reduced dependency on codon optimality ^48^ and coupling of 3’UTR translation to piRNA biogenesis ^49^. In this context, heterogenous ribosome dwell times at stop codons may facilitate functional partitioning of ribosomes beyond annotated coding regions. Whether termination dynamics are developmentally regulated during spermatogenesis remains an important question for future study.

To identify cell type-specific regulators of termination pausing, we focused on the eukaryotic-specific ribosomal protein Rps26. Like the 3’ end of 18S rRNA, Rps26 crosslinks to mRNA regions upstream of the P-site ^31,37^, and occupies a strategic position beneath the mRNA segment upstream of the E-site, effectively shielding it from direct interaction with 18S rRNA (Figure 7A). Loss of Rps26 enhances termination pausing at GA-rich stop codons, whereas Rps26 overexpression dampens pausing, consistent with a model in which Rps26 restricts mRNA:rRNA base pairing. Intriguingly, Rps26 was previously shown to regulate initiation by recognizing Kozak sequence elements ^50^. The consensus vertebrate Kozak context (GCCRCC<u>AUG</u>G) highlights the importance of purine at key positions, paralleling our observations at stop codons. The reciprocal changes in initiation and termination peaks upon Rps26 depletion suggest that coordinated mRNA:rRNA:Rps26 interactions operate throughout the translation cycle.

Perhaps the most expected finding of our study is that tissue-specific termination pausing is governed by Rps26 stoichiometry. In yeast, Rps26 can dissociate from fully assembled 80S ribosomes under stress ^38^, raising the possibility that mammalian ribosomes similarly exist in Rps26-variable states. is either sub-stoichiometric or loosely integrated into the 80S ribosomes. Notably, Rps26 haploinsufficiency is linked to Diamond-Blackfan anemia (DBA) ^51^. Prolonged termination pausing and impaired ribosome recycling may contribute to the hematologic defects observed in DBA, particularly given the high reliance of erythroid cells on ribosome rescue pathways and their elevated levels of 3’UTR translation ^52^. A deeper understanding of stop codon-associated translational quality control may therefore open new avenues for therapeutic interventions in DBA and other diseases caused by premature termination of nonsense mutations.

## ACKNOWLEDGMENTS

We thank Cornell University Life Sciences Core Laboratory Center for sequencing and FACS. S.U. was supported by Takeda Science Foundation. This work was supported by US National Institutes of Health (DP1GM142101) and HHMI Faculty Scholar (55108556) to S.-B.Q.

## AUTHOR CONTRIBUTIONS

S.-B.Q. conceived the project. L.J. performed most experiments. Y.M. conducted the majority of sequencing data analysis. S.U. performed data analysis involving 18S rRNA and Rps26. A.X.L. performed tissue polysome analysis of Rps26. L.D. contributed to the sequencing of mouse tissues. L.H.F.L. conducted ribosome structural analysis. S.-B.Q. wrote the manuscript. All authors initially discussed the results and edited the manuscript.

## STAR*METHODS

## KEY RESOURCES TABLE

| REAGENT or RESOURCE | SOURCE | IDENTIFIER |
| --- | --- | --- |
| **Antibodies** |  |  |
| Anti-GFP antibody | Proteintech | Cat# 50430-2-AP; RRID: AB_11042881 |
| Anti-eRF1 antibody | Santa Cruz Biotechnology | Cat# sc-365686; RRID: AB_10843214 |
| Anti-eRF3 antibody | Santa Cruz Biotechnology | Cat# sc-515615 |
| Anti-Rps26 antibody | Proteintech | Cat# 14909-1-AP; RRID: AB_2180361 |
| Anti-Rpl4 antibody | Proteintech | Cat# 11302-1-AP; RRID: AB_2181909 |
| Anti-ABCE1 antibody | Abcam | Cat# ab185548; RRID: AB_2858278 |
| Anti-myc antibody | Cell Signaling | Cat# 2272S |
| Anti-RACK1 antibody | BD Transduction Laboratories | Cat# 610177; RRID: AB_397576 |
| Anti-β-Actin antibody | Millipore Sigma | Cat# A5441; RRID: AB_476744 |
| Anti-Mouse IgG (Fc specific)-Peroxidase antibody | Millipore Sigma | Cat# A0168; RRID: AB_257867 |
| Anti-Rabbit IgG (whole molecule)-Peroxidase antibody | Millipore Sigma | Cat# A9169; RRID: AB_258434 |
| **Bacterial and virus strains** |  |  |
| DECIPHER pRSI9-U6-(sh)-UbiC-TagRFP-2A-Puro | Cellecta | N/A |
| Subcloning Efficiency DH5a Competent Cells | Thermo Fisher Scientific | Cat# 18265-017 |
| **Chemicals, peptides, and recombinant proteins** |  |  |
| Dulbecco's Modified Eagle Medium | Corning | Cat# 10-013-CV |
| Fetal bovine serum | Millipore Sigma | Cat# 12306C-500ML |
| Opti-MEM I Reduced Serum Medium | Thermo Fisher Scientific | Cat# 31985-070 |
| Dulbecco's Phosphate Buffered Saline | Thermo Fisher Scientific | Cat# 14190250 |
| BamHI-HF | New England BioLabs | Cat# R3136S |
| HindIII-HF | New England BioLabs | Cat# R3104S |
| Lipofectamine MessengerMAX | Invitrogen | Cat# LMRNA015 |
| Lipofectamine 2000 | Invitrogen | Cat# 11668-500 |
| DL-Dithiothreitol | Millipore Sigma | Cat# D0632-5G |
| Blotting-Grade Blocker | Bio-Rad | Cat# 1706404 |
| Tween-20 | Millipore Sigma | Cat# P7949-500ML |
| Triton-X100 | Millipore Sigma | Cat# T9284-100mL |
| EDTA-free Protease Inhibitor Cocktail Tablets | Millipore Sigma | Cat# 11 836 170 001 |
| Puromycin | Millipore Sigma | Cat# P7255 |
| Cycloheximide | Millipore Sigma | Cat# C1988-1G |
| TRIzol LS Reagent | Invitrogen | Cat# 10296-028 |
| Nuclease-Free Water | Invitrogen | Cat# AM9932 |
| SUPERase-In RNase Inhibitor | Invitrogen | Cat# AM2696 |
| RNase I | Invitrogen | Cat# AM2295 |
| 15% TBE-Urea Gels | Invitrogen | Cat# EC6885BOX |
| SYBR Gold nucleic acid gel stain | Invitrogen | Cat# S-11494 |
| Sodium acetate buffer solution | Millipore Sigma | Cat# S7899-500ML |
| T4 Polynucleotide Kinase | New England BioLabs | Cat# M0201L |
| E. coli Poly(A) Polymerase | New England BioLabs | Cat# M0276L |
| T4 RNA Ligase 2, truncated | New England BioLabs | Cat# M0242L |
| Ezra enzyme | This paper | N/A |
| UltraPure™ SSC, 20X | Invitrogen | Cat# 15557044 |
| Streptavidin Magnetic Beads | New England BioLabs | Cat# S1420S |
| RNaseOUT Recombinant Ribonuclease Inhibitor | Invitrogen | Cat# 10777-019 |
| 8% TBE Gel | Invitrogen | Cat# EC6215BOX |
| Pierce Protein A/G Agarose | Thermo Fisher Scientific | Cat# 20421 |
| Critical commercial assays |  |  |
| QIAquick Gel Extraction Kit | Qiagen | Cat# 28706 |
| Q5 Site-Directed Mutagenesis Kit | New England BioLabs | Cat# E0554S |
| mMESSAGE mMACHINE T7 Transcription Kit | Invitrogen | Cat# AM1344 |
| Poly(A) Tailing Kit | Invitrogen | Cat# AM1350 |
| RNA Clean and Concentrator-25 Kit | Zymo | Cat# R1018 |
| Nano-Glo HiBiT Lytic Detection System | Promega | Cat# N3040 |
| Nano-Glo HiBiT Blotting System | Promega | Cat# N2410 |
| SuperScript III Reverse Transcriptase | Thermo Fisher Scientific | Cat#18080-044 |
| Deposited data |  |  |
| Raw and analyzed data | This paper | GEO: |
| Experimental models: Cell lines |  |  |
| Human: HEK293-K<sup>b</sup> | Laboratory of Jonathan Yewdell | N/A |
| Human: Lenti-X 293T | Takara Bio | Cat# 632180 |
| Experimental models: Organisms/strains |  |  |
| Mouse: C57BL/6 | Jackson Laboratory | N/A |
| Oligonucleotides |  |  |
| Oligonucleotide sequences used in this study are provided in Table S1 | This paper | N/A |
| Recombinant DNA |  |  |
| pcDNA3.1 (myc-His B)-Rps26 | This paper | N/A |
| pRL-CMV-18S rRNA wild type | Burman and Mauro, 2012 | N/A |
| pRL-CMV-18S rRNA mutant | This paper | N/A |
| Software and algorithms |  |  |
| GraphPad Prism | Dotmatics | RRID: SCR_002798 |
| Snapgene | Dotmatics | RRID: SCR_015052 |
| R | The R Project | https://www.r-project.org/ |
| Custom Code | This paper |  |

## RESOURCE AVAILABILITY

### Lead contact

All material request should be directed to Shu-Bing Qian.

### Materials availability

Reagents and materials produced in this study are available from the Lead Contact pending a completed Materials Transfer Agreement.

### Data and code availability

- All Sequencing data are available in the Gene Expression Omnibus database. All raw images are available in Mendeley Data. All data are publicly available as of the date of publication. Accession numbers and DOI are listed in the key resources table.
- All custom code has been deposited to GitHub and Zenodo. DOI are listed in the key resources table.
- Any additional information required to reanalyze the data reported in this paper is available from the lead contact upon request.

## EXPERIMENTAL MODEL AND SUBJECT DETAILS

### Cell lines

HEK293-K^b^ cells and Lenti-X 293T cells are maintained in Dulbecco’s Modification of Eagle’s Medium (Corning, 10-013-CV) with 10% fetal bovine serum (Sigma, 12306C). All cells were grown at 37°C with 5% CO_2_.

### Mouse strains

C57BL/6 mice were obtained from the Jackson laboratory. All animals (1-6 mice per cage) were housed in a 12 h light/dark cycle in the Weill Hall animal facility at Cornell University with the supervision of the Center for Animal Resources and Education (CARE) breeding program. All animals used in this study were handled in accordance with federal and institutional guidelines, under a protocol approved by the Cornell University Institutional Animal Care and Use Committee, protocol 2017-0035. Mice were housed under specific pathogen-free conditions in an Association for the Assessment and Accreditation of Laboratory Animal Care International-accredited facility and cared for in compliance with the Guide for the Care and Use of Laboratory Animals.

## METHOD DETAILS

### Antibodies

The following antibodies were used at their indicated experimental concentrations: anti-GFP (Proteintech, 50430-2-AP, 1:1000), anti-eRF1 (Santa Cruz Biotechnology, sc-365686, 1:200), anti-eRF3 (Santa Cruz Biotechnology, sc-515615, 1:200), anti-Rps26 (Proteintech, 14909-1-AP, 1:500), anti-Rpl4 (Proteintech, 11302-1-AP, 1:1000), anti-ABCE1 (Abcam, ab185548, 1:1000), anti-myc (Cell Signaling, 2272S, 1:1000), anti-RACK1 (BD Transduction Laboratories, 610177, 1:1000) and anti-β-Actin ((Sigma-Aldrich, A5441, 1:5,000). anti-mouse IgG horseradish peroxidase (HRP)-conjugated secondary antibodi (Sigma-Aldrich, A0168, 1:10,000) or anti-rabbit IgG secondary antibody conjugated to peroxidase (Sigma-Aldrich, A9169, 1:10,000).

### Plasmid construction

The template of EGFP-HiBiT reporters were PCR-amplified from the pcDNA3-EGFP vectors using reverse primer containing the desired sequence. In some cases, the template of reporters based on HiBiT-EGFP were generated using a two-step PCR amplification approach. First, the full length of HiBiT and EGFP was amplified from pcDNA3-EGFP to generate HiBiT-EGFP. The resulting PCR product was used as a template to produce the full-length reporter using a second forward primer containing the respective sequence. After column purification (QIAGEN), the DNA template (1∼2 μg) was utilized to generate mRNAs suitable for transfection. For exogenous Rps26, the full-length coding sequence of human Rps26 was cloned into pcDNA3.1(myc-His B) using BamH I and Hind III restriction sites. To create the 18S rRNA mutant, site-directed mutagenesis was performed using Q5 Site-Directed Mutagenesis Kit (New England Biolabs) according to the manufacturer manual. Mutation was confirmed by Sanger DNA sequencing. DNA sequences of all primers used in this study are listed in the Key Resources Table.

### In vitro transcription

To prepare mRNA reporters, 1∼2 μg PCR products described above were utilized as templates to generate mRNAs suitable for transfection. In vitro transcription was performed for 1 h at 37 °C using mMESSAGE mMACHINE T7 Transcription Kit (Invitrogen) followed by poly(A) tailing (Ambion) at 37 °C for 30 min. The resulting RNAs were purified using RNA Clean & Concentrator (Zymo Research) following the manufacturer’s instruction and stored at -80 °C.

### Transfection

For mRNA reporter transfection, cells were transfected with in vitro transcribed mRNA (1 μg) in Opti-MEM (125 µl) using Lipofectamine MessengerMAX (1 µl) in Opti-MEM (125 µl), unless stated otherwise. The cells were incubated with the mRNA/Lipofectamine MessengerMAX mixture for 4 h followed by immunoblotting, HiBiT assay or polysome profiling. For Rps26 or 18S rRNA overexpression, 2 μg plasmids were mixed with 4 μl Lipofectamine 2000 (Invitrogen) followed by incubation with cells for at least 24 h, unless stated otherwise.

### HiBiT assay

Cells grown in a 35 mm dish were transfected with mRNA reporters (1 μg) described above. Transfected cells were washed with PBS and then lysed using a Nano-Glo HiBiT Lytic Detection System (Promega) according to manufacturer’s instructions. HiBiT signals were measured using Luminometer (Atto).

### Immunoblotting

Cells grown in a 6-well plate were transfected with transcribed mRNA reporters (1 μg) described above. Transfected cells were washed twice with ice-cold PBS and lysed on ice by adding SDS-PAGE sample buffer (50 mM Tris pH 6.8, 100 mM DTT, 2% SDS, 0.1% bromophenol blue, 10% glycerol), followed by heating for 10 min at 95 °C. Protein samples were separated on SDS-PAGE gels followed by transferring to PVDF membranes (Thermo Fisher Scientific). Membranes were blocked in 5% non-fat milk (Bio-Rad) in TBS containing 0.1 % Tween-20 (TBST) for 1 h, followed by incubation overnight with primary antibodies at 4 °C. After 3 × 10 min washes in TBST, membranes were incubated with anti-mouse IgG horseradish peroxidase (HRP)-conjugated secondary antibodies at room temperature for 1 h. Membranes were then washed 3 × 10 min in TBST at room temperature and visualized using chemiluminescence by exposing to ECL film (GE Healthcare).

To prepare tissue lysates, mouse tissues were dissected and snap-frozen in liquid nitrogen. Frozen tissues were thawed and homogenized on ice with homogenizer (U.S. Solid) in ice-cold RIPA buffer (150 mM NaCl, 50 mM Tris-HCl pH 8.0, 1% Triton X-100, 0.5% sodium deoxycholate, 0.1% SDS) with 1 × protease inhibitors (Roche). After centrifugation at 16,000 g for 20 min at 4 °C, supernatants were collected for immunoblotting as described above. Proteins in mouse tissue fractions collected from sucrose density gradients were precipitated by trichloroacetic acid (TCA) and resuspended in 2 × SDS-containing sample buffer. To detect HiBiT-tagged proteins by Nano-Glo HiBiT blotting system (Promega), cell lysates were resolved by SDS-PAGE and transferred to PVDF membranes as described above. The membrane was incubated with 1 × TBST for 1 h at room temperature, followed by incubation in the LgBiT/buffer solution (50 μl of LgBiT protein in 10 ml of Nano-Glo blotting buffer) at room temperature for 1 h. 20 μl of substrate was added to the incubation solution for additional 5 min. The membrane was exposed to ECL film in the same manner as the immunoblot analysis.

### shRNA knockdown

shRNAs targeting Rps26 and ABCE1 were designed from BROAD RNAi consortium database and subcloned into DECIPHER pRSI9-U6- (sh)-UbiC-TagRFP-2A-Puro (Cellecta). A scrambled shRNA was used as control. Lentiviral particles were produced using Lenti-X 293T cells (Clontech). The supernatants containing viral particles were harvested at 48 h after transfection and filtered through a 0.45 μM Millex-HP filter unit (Millipore). HEK293 cells were transduced with shRNA lentivirus for 48 h followed by selection with 2 µg/ml puromycin. Knockdown efficiency was detected by immunoblotting using indicated antibodies. The oligonucleotide sequences are listed in the Key Resources Table.

### Polysome profiling

A total of 4 plates (10-cm) HEK293 cells grown to 80% confluency were washed by cold PBS and lysed in the polysome lysis buffer (10 mM HEPES, pH 7.4, 100 mM KCl, 5 mM MgCl_2_, 100 μg/mL cycloheximide with 1% Triton X-100). The nuclei were pelleted by spinning at 14,000 rpm for 10 min at 4 °C. For mouse tissues, 100 mg of frozen samples were homogenized on ice using a Dounce homogenizer in 1 mL polysome lysis buffer. Homogeneous lysates were cleared by centrifugation at 14,000 rpm for 10 min at 4 °C. 500 µL of lysates were loaded onto a 15-45% (wt/vol) sucrose density gradients freshly prepared in a SW41 ultracentrifuge tube (Backman) using a Gradient Master (BioComp Instruments). Samples were centrifuged at 180,000 g for 2 h 30 min at 4 °C in a Beckman SW41 rotor. Polysome profiles were recorded at A254 using the Brandel Gradient Fractionation System and an ISCO UA-6 UV/Vis detector.

### Ribosome profiling

The Ezra-seq has been described previously ^22^. In brief, an aliquot of ribosome fractions representing monosome or polysome were collected followed by digestion with *E. coli* RNase I (Ambion, 750 U per 100 A260 units) by incubation at 4 °C for 1 h. RNA was extracted using Trizol LS reagent (Invitrogen) followed by ethanol precipitation. The ribosome-protected mRNA fragments (RPFs) were separated on a 15% polyacrylamide TBE-urea gel (Invitrogen) and visualized using SYBR Gold (Invitrogen). Selected regions in the gel corresponding to 25-35 nt were excised and dissolved by soaking in 400 μl RNA elution buffer (300 mM NaOAc pH 5.2, 1 mM EDTA, 0.1 U/μl SUPERase·In) for 10 min at 70 °C. The gel debris was removed using a Spin-X column (Corning), followed by ethanol precipitation. 14 μl RNAs (10∼200 ng) were mixed with 1 μl T4 PNK (NEB), 20 U SUPERase·In in 1 × T4 PNK buffer and incubated at 37 °C for 30 min followed by 65 °C for 20 min. After ethanol precipitation, 10 μl dissolved RNA were mixed with 1 μl homemade Ezra enzyme, 1 μl Poly(A) Polymerase (NEB) and 20 U SUPERase·In in 7 μl Ezra buffer. After incubation at 37 °C for 30 min followed by 65 °C for 20 min, 1 μl of 1 μM 5’ end adaptor, 1 μl of 1 μM biotinylated reverse transcription primer, 20 U SUPERase·In were added and incubated at 70 °C for 3 min followed by slowly cooling down (3 °C/min) to 25 °C. The hybridized RNA sample was mixed with 10 μl of pre-washed streptavidin beads and incubated at room temperature for 10 min. Beads were washed and re-suspended in 10 μl nuclease-free water. Ligation was performed for 60 min at 25 °C by mixing beads with a 10 μl reaction mixture (1 × T4 Rnl2 reaction buffer, 20 U SUPERase·In, 15% PEG8000 and 200 U T4 RNA ligase 2 truncated KQ (NEB)). After washing once with 2 × SSC, beads were re-suspended in 12 μl nuclease-free water and mixed with 8 μl cDNA synthesis mixture (5 × first strand buffer, 0.1 M DTT, 10 mM dNTP, RNaseOUT and SuperScript III) followed by incubation at 50 °C for 30 min. After washing once with 2 × SSC, beads were resuspended in 10 μl nuclease-free water and incubated at 95 °C for 2 min, then immediately placed on the ice for 1 min. After placing on magnet stand for 1 min, the supernatant cDNA was amplified by PCR using barcoded sequencing primers. PCR was performed by mixing 1 × HF buffer, 0.5 mM dNTP, 0.25 μM PCR primers and 0.025 U Phusion polymerase. PCR was carried out under the following conditions: 98 °C, 30 s; (98 °C, 5 s; 68 °C, 15 s; 72 °C, 10 s) for 12 cycles; 72 °C, 3 min. PCR products were separated on a 8% polyacrylamide TBE gel (Invitrogen). DNA products with the expected size 180 bp were excised and recovered from DNA elution buffer (300 mM NaCl, 1 mM EDTA). After quantification by Agilent BioAnalyzer DNA 1000 assay, equal amounts of barcoded samples were pooled and sequenced using NextSeq 500 (Illumina). The oligonucleotide sequences are listed in the Key Resources Table.

### eRF1-seq

A total of 4 plates (10-cm) HEK293 cells with 80% confluence were washed three times with ice-cold DPBS. Cells were fixed in ice-cold formaldehyde solution (0.5% in DPBS) for 10 min at 4 °C on a rocker. After washing with ice-cold DPBS three times, cells were quenched in ice-cold buffer (50 mM Glycine, 50 mM Tris-HCl pH 7.5 in nuclease-free water) for 10 min at 4 °C on a rocker. Cells were then washed with polysome buffer and lysed in the 400 μl of polysome lysis buffer with 1% Triton-X-100 on ice. Cell debris was removed by centrifugation at 15,000 rpm for 10 min at 4 °C. The supernatant was digested with RNase I (Ambion, 750 U per 100 A260 units) for 1 h at 4 °C. Digested supernatant was loaded onto sucrose gradients for polysome profiling as described above. The 80S fraction was collected (∼200 μl total) and mixed with 10 μg eRF1 antibody and 0.5 U/μl SUPERase·In (Invitrogen) followed by incubation under gentle rotation at 4 °C for 5 h. Protein A/G beads were added into the mixture and rotated at 4 °C overnight. Beads were washed three times and then resuspended in 600 μl of polysome buffer. RNA was extracted from resuspended beads in polysome buffer. Briefly, samples were brought to room temperature and then adjusted to 10 mM Tris-HCl pH 7.4, 10 mM glycine, 1% (w/v) SDS and 10 mM EDTA pH 8.0 followed by incubation at 65 °C for 5 min. One volume of acidic phenol/chloroform solution was added and vortexed at maximum speed for 2 min. Mixtures were then placed into thermomixer and shake at 1,400 rpm for 20 min at 65 °C to reverse the cross-links. After centrifugation at 14,000 rpm for 5 min at 4 °C, the aqueous phase was precipitated with ethanol. Purified RNA was used for cDNA library construction and high-throughput sequencing as described above.

### Massively paralleled reporter assay (MPRA)

From the PCR product of HiBiT-EGFP described above, a second PCR was conducted using the pooled oligonucleotide library (IDT) and a primer containing the T7 promoter. The DNA template (1∼2 μg) was utilized to generate the mRNA library via in vitro transcription as described above. Cells with 80% confluence were transfected with 6 µg of mRNA library using Lipofectamine MessengerMAX. Cells were lysed 4 h after transfection followed by polysome profiling as described above. Fractions of 500 µl corresponding to monosome or polysome were collected for RNA extraction using TRIzol LS. RNA was purified using RNA Clean & Concentrator and eluted with 11 µl of nuclease-free water. The purified RNA was reverse transcribed using SuperScript III and gene-specific primers. In brief, RNA samples were mixed with 1 μl of 10 mM dNTP, 2 pmol reverse primer and incubated at 65 °C for 5 min, then immediately placed on ice for 1 min. The reverse transcription was carried out by incubating with the 7 μl reaction mixture (5 × first strand buffer, 0.1 M DTT, RNaseOUT and SuperScript III) at 50 °C for 60 min followed by heating at 70 °C for 15 minutes. The products were then amplified with Illumina-based sequencing primers with barcode. PCR were performed by mixing 1 × HF buffer, 0.5 mM dNTP, 0.25 μM PCR primers and 0.025 U Phusion polymerase. The PCR was initiated at 98 °C, 30 s; then (98 °C, 5 s; 68 °C, 15 s; 72 °C, 10 s) for 12 cycles; 72 °C, 3 min. The PCR products with the expected size 190 bp were excised from a 8% polyacrylamide TBE gel. The DNA products were recovered from DNA elution buffer, followed by quantification using Agilent BioAnalyzer DNA 1000 assay. Equal amounts of barcoded samples were pooled for sequencing using NextSeq 500 (Illumina). The oligonucleotide sequences are listed in the Key Resources Table.

### Structural analysis

To gain insights about the relative orientations and interaction likelihood between the 5’ end of the mRNA (site -13 to -3 related to the A-site) and the 3’ end of the 18S rRNA along the ribosome translocation, a structural analysis was carried using the respective PDBs 6GZ3 (with a 3.60 Ǻ resolution), 6GZ4 (3.60 Å), 6GZ5 (3.50 Å) and 6yal (3.00 Å). The PDBs 6GZ3, 6GZ4 and 6GZ5 encompass three respective intermediate snapshots between the PRE and the POST translocation steps for the eukaryotic 80S ribosome (hereafter, referred simply as PRE and POST states). These three structures (hereafter referred as translocation intermediates POST 1 to 3, or simply TI-POST 1 – 3 for the respective PDBs 6GZ3 – 5) were solved, described and discussed previously ^33^. The mRNA environment sequentially described by these three PDBs provides a reasonable glimpse about the changes occurring after the recognition of at the A site codon and along the movement of the ribosome to the next codon. The PDB 6YAL, in turn, is the Homo sapiens 48S late-stage initiation complex ^53^. Due to the higher resolution of the PDB 6YAL, its mRNA structure is solved until a higher 5’ extension compared to the 6GZ3–5 PDBs (The mRNA 5’ in 6YAL starts from the -18 nt related to the site A, while in 6GZ3 it starts from the site -7 and from the site -6 in 6GZ4 and 6GZ5). In this way, we have made use of the 5’ fragment of the 6YAL mRNA structure to build a rigid body model of the same extension starting from the -13 nt at each one of the PDBs 6GZ3–5, using structural alignment of the respective backbone atoms in each oligonucleotide extremity in Pymol [Schrodinger, LLC. 2010. The PyMOL Molecular Graphics System, Version 2.5].

### Anisotropic network modeling

The relative local fluctuations of the exit mRNA channel at the ribosome structure and the consequent proximity likelihood between the mRNA 5’ extension and the rRNA 3’ end was estimated in presence and absence of Rps26 by normal mode analysis (NMA) using anisotropic network modeling (ANM) ^54^. For this analysis, we used the atoms from the PDB 6YAL around a 30 Ǻ region centered on Rps26 (depicting the mRNA channel exit) both in the presence and absence of this protein. The PDB 6YAL was chosen due to its higher resolution as a whole, besides longer extension of the mRNA 5’ end compared to the other three PDBs structurally analyzed in this study. Furthermore, the region around 30 Ǻ from Rps26 (hereafter called Rps26 site) encompass a symmetric sphere composed basically of residues from the 40S subunit (including the 18S 3’ extension), mRNA (sites -13 to +5 nt related to the A-site) and the anti-codon loop from the P-site tRNA, common to all the four structures here analyzed. Finally, this region in 6YAL presents a relatively small global root mean square deviation (RMSD), considering the protein and RNA backbone, related to both PDBs 6GZ3 and 6GZ5 (1.478 Å in the two cases). In this way, the PDB 6YAL was considered an accurate approximation of the general environment of the Rps26/mRNA 5’ end/rRNA 3’ end triad in ribosome, despite portraying a pre-initiation structure.

The ANM was carried using the ProDy tools ^55–57^. A Hessian matrix was built upon the backbone atoms of the Rps26 site for both proteins (Cα) and RNA (P, C4’ and C2) with and without Rps26 (hereafter referred as + Rps26 and – Rps26). The mRNA was considered from the -10 to the +5 nt, once the first three residues (-13 to -11) are more distant from the rRNA 3’ end and free of direct contacts with the neighborhood as a whole, which makes their movements dominate the NMA if they are considered on the ANM (not shown). The Hessian matrix was configured using the default parameters of distance cutoff and gamma function, with the respective values of 15 Å and 1.0 kcal/(mol.Å2). Initially, the first 50 normal modes of each system were estimated by obtaining their respective covariance matrixes by diagonalizing the Hessiam matrixes. The modes simultaneously containing the largest possible eigenvalues and higher mRNA fluctuations at the -10 to -3 extension (directly parallel to the rRNA 3’ end and separated from it by Rps26), as well as the fluctuations of the last 10 residues at the 18S 3’ end, were selected. In this way, the 15 first ANM normal modes estimated from the + Rps26 and – Rps26 systems were taken for conformational sampling analysis.

The +Rps26 and –Rps26 structures of the Rps26 site from above were taken to sampling of alternate conformations along the global fluctuations described by their respective 15 first ANM modes using ProDy ^55–57^. Basically, an extended ANM model containing all the protein and rRNA atoms for each structure was built from the original coarse grain model containing only the backbone atoms used to build the Hessian matrix. All the atoms at the extended model still obey the movements dictated by the selected 15 first normal modes, with each side chain atom moving in the same direction that the backbone atoms of the residue to which they belong. A set of 70 conformations symmetrically distributed along the fluctuation governed by the 15 first normal modes and with an average RMSD of 2.5 Å related to the input structure was sampled for each one of the +Rps26 and –Rps26 models. Finally, from each original 70 conformers set, a subset containing only the sterically feasible structures (i.e., without significant clashes or conformational distortions) was taken for analysis. Although higher refinements would be necessary to take these final conformers to rigorous molecular dynamics or free energy calculation studies, they provide enough insights about the Rps26 influence on the mRNA 5’ end/18S 3’ end fluctuations and interaction distance likelihood at the ribosome context.

## QUANTIFICATION AND STATISTICAL ANALYSIS

Data is presented as mean ± SEM, unless otherwise stated. At least three independent biological replicates have been performed for each experiment. The number of independent experiments is indicated. Statistical tests used and specific p-values are indicated in the figure legends.

### Analysis of Ribo-seq and eRF1-seq

The adaptor of sequencing reads was clipped by Cutadapt, using parameters: -a AAAAAA --max-n=0.1 -m 15. The clean reads were then aligned to human transcriptome (GRCh38.81), which contains the protein coding transcripts with the longest CDS, using STAR with default parameters. To avoid ambiguity, reads mapped to multiple positions or with > 2 mismatches were disregarded for further analysis. Ribosome P-site was defined as the positions of 12th, 13th and 14th from 5’ end of the read (position 0). A-site was defined as the positions of 15th, 16th and 17th. To generate aggregation plot around the start and stop codons, for each mRNA, the aligned reads at individual sites were normalized by mean reads of the CDS. mRNAs with total reads in CDS < 16 or the CDS sites covered by footprints < 10% were excluded. The normalized values of the sites with the same distance relative to the start codon or stop codon were averaged across transcriptome.

### Identification of termination peaks

To identify termination peaks, all reads of eRF1-seq were assigned to individual sites on mRNAs. The mRNAs with < 10 total reads from eRF1-seq were excluded. A 120-nt sliding window was used to scan along the mRNA, the sites with terminating reads tenfold higher than the average reads within the sliding window were defined as the termination peaks.

### uORF prediction

For each mRNA, all possible uORFs starting with AUG were first extracted. A Wilcoxon test was applied to test whether the in-frame reads are significantly higher than the other two frames. The two *P* values were then combined to a single *P* value using a Stouffer’s method. uORFs with a false discovery rate (FDR) <0.05 were defined as the uORFs with robust translation.

### RNA secondary structure analysis

A 30-nt sliding window was used to scan 3’ UTR. For each window, the minimum fold free energy (MFE) was calculated by ViennaRNA [PMID: 22115189] using default parameters.

### Analysis of MPRA dataset

For each raw sequencing file, the adaptors at both ends were removed by cutadapt. The trimmed reads with length unequal to 9 nucleotides were excluded from analysis. The remaining trimmed reads were counted and then an RPM value (reads per million) was obtained by dividing the resultant read count by the total count.

