## Supplementary Fig 1 - 7 for "Profiling of terminating ribosomes reveals translational control at stop codons"

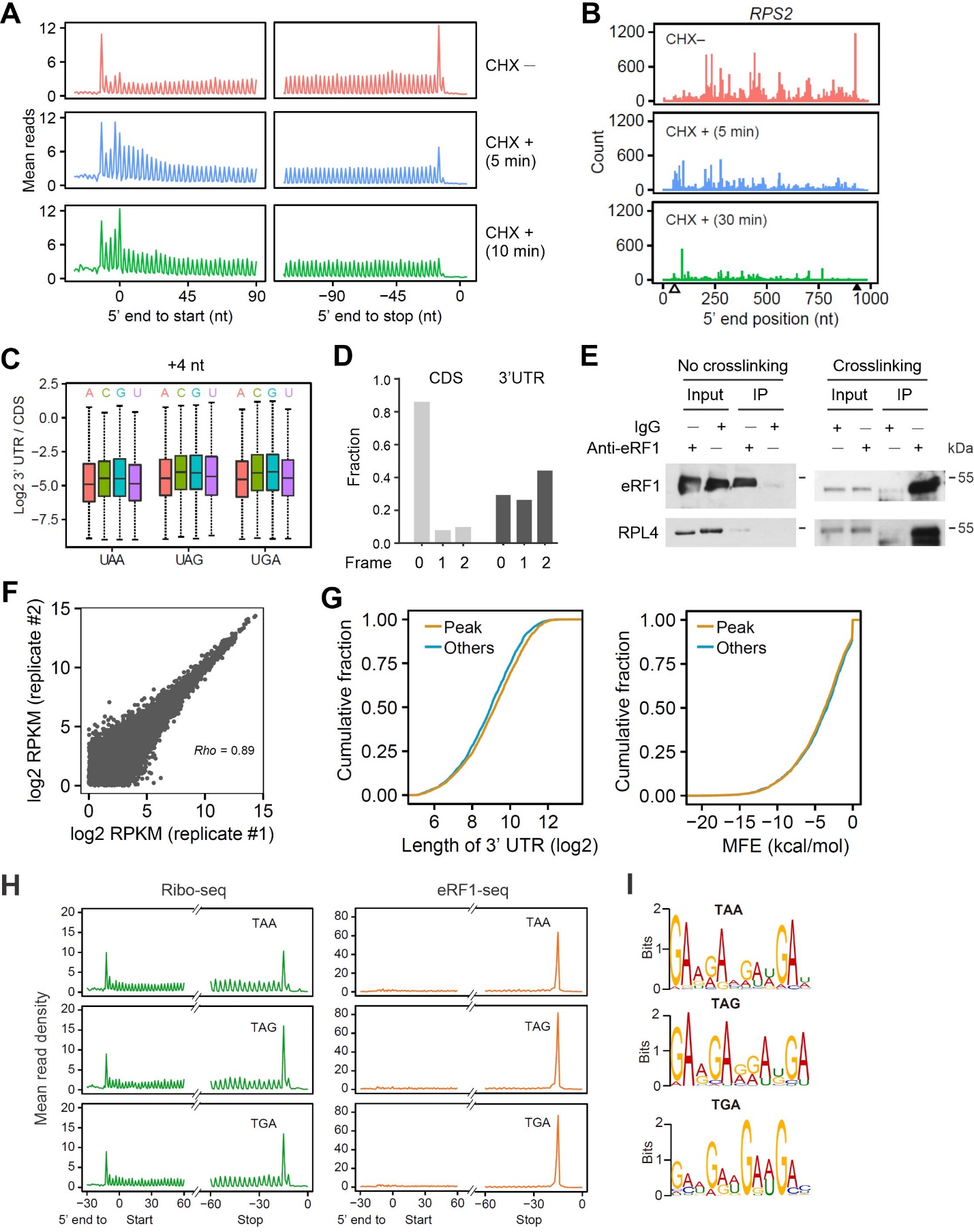

**Figure S1. Characterizing terminating ribosomes.**

(A) HEK293 cells were treated with cycloheximide (CHX, 100 μg/mL) for varied times followed by Ribo-seq. Aggregation plots show the distribution of mean reads across the transcriptome aligned at start and stop codons.
(B) A representative mRNA (*RPS2*) with reads obtained from ribo-seq in cells as (A). The 5’ end of reads were used for mapping.
(C) The ratio of ribosome density in 3’ UTR over the density in CDS. The mRNAs were stratified into different groups based on the identify of stop codons and the nucleotide immediately after the stop codon.
(D) Relative fractions of Ribo-seq reads mapped to different reading frames. Reads in CDS and 3’UTR were shown.
(E) HEK293 cells with or without crosslinking by formaldehyde were subjected to eRF1 immunoprecipitation followed by Western blotting using antibodies indicated. Representative results of three independent experiments are shown.
(F) A scatter plot shows the correlation of read counts between two biological replicates of eRF1-seq (*Rho* = 0.89).
(G) Comparison of 3’UTR length (left) and folding free energy (right) between mRNAs with and without eRF1 peaks at the stop codon.
(H) Aggregation plots of ribosome density using Ribo-seq and eRF1-seq data sets on mRNAs with different stop codons.
(I) The enriched sequence motif preceding different stop codons on mRNAs with eRF1 peaks.

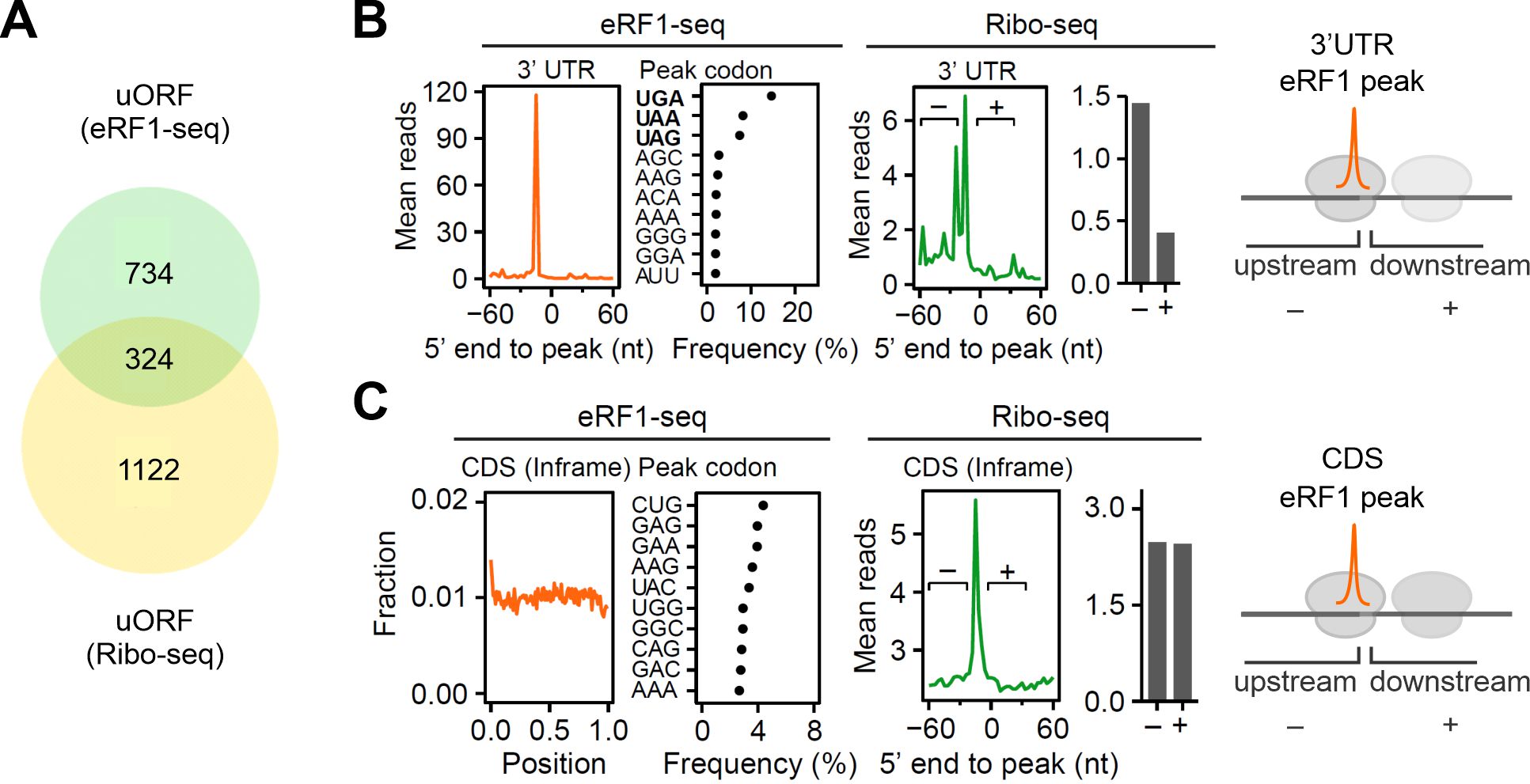

**Figure S2. eRF1-seq reveals prevailing termination sites in 3’UTR and CDS.**

(A) A Venn diagram shows the number of uORFs identified by Ribo-seq and eRF1-seq.
(B) The left panel shows mean eRF1-seq reads around the position of eRF1 peaks within 3’ UTR. The dot plot shows the frequency of A-site codons at eRF1 peaks. The right panel shows mean Ribo-seq reads around the position of eRF1 peaks within 3’ UTR. The bar graph shows the mean ribosome densities before (-) and after (+) the eRF1 peaks.
(C) The left panel shows mean eRF1-seq reads around the position of in-frame eRF1 peaks within CDS. The dot plot shows the frequency of A-site codons at eRF1 peaks. The right panel shows mean Ribo-seq reads around the position of in-frame eRF1 peaks within CDS. The bar graph shows the mean ribosome densities before (-) and after (+) the eRF1 peaks.

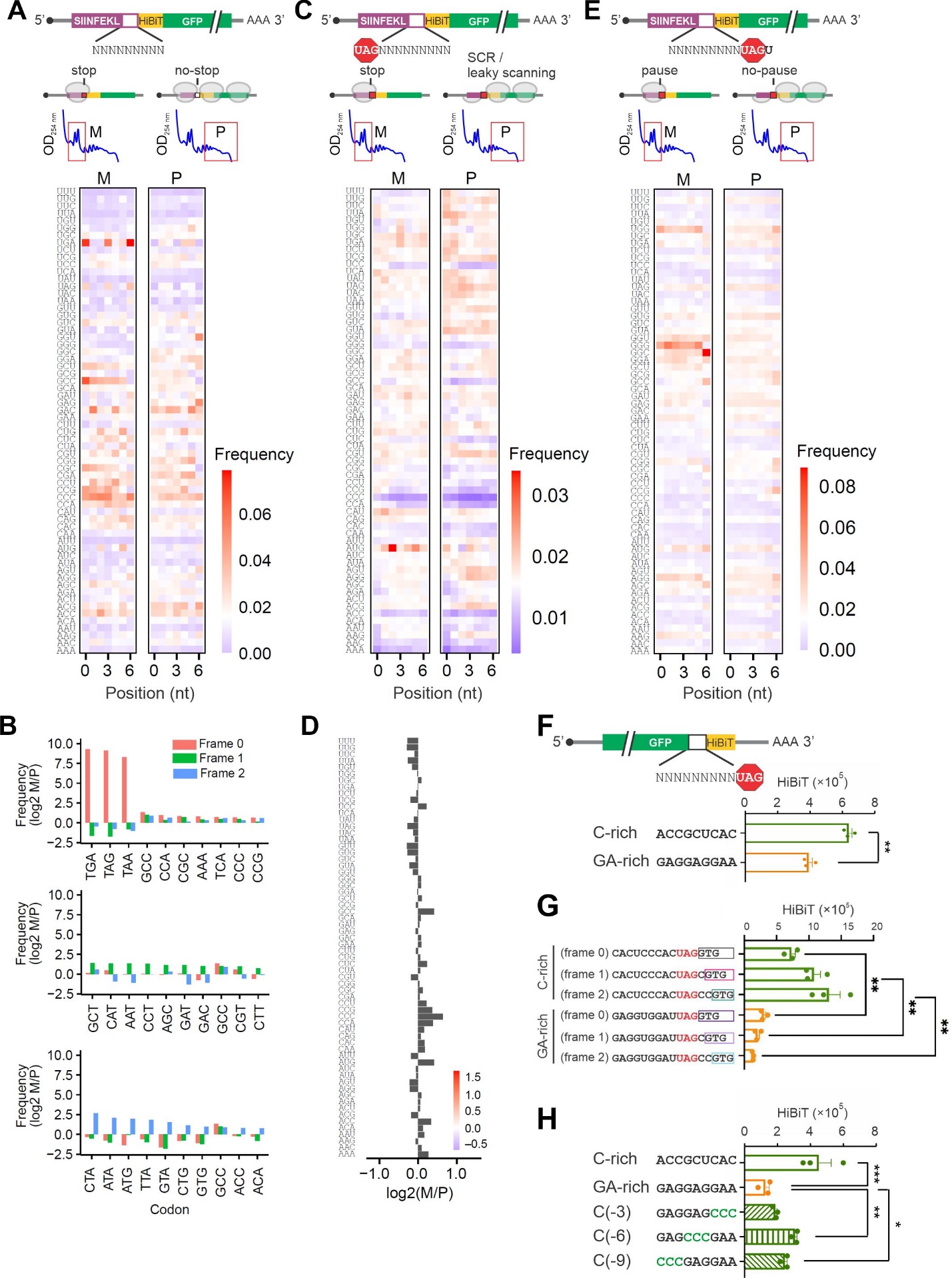

**Figure S3. MPRA assays dissect the sequence context of stop codons.**

(A) The up panel shows the schematic of a massively parallel reporter assay, where the stop codon of uORF was replaced with a random 9-nt sequences. The uORF translation was monitored by the number of associated ribosomes separated by sucrose gradient (M, monosome; P, polysome). The heatmap shows the codon frequency within the 9-nt random sequence in monosome (M) and polysome (P) fractions.
(B) Bar graphs show the top 10 codons enriched in monosome (M) over polysome (P) at each reading frame.
(C) The up panel shows the schematic of a massively parallel reporter assay, where a random 9-nt sequence was replaced after the uORF stop codon UAG. The uORF translation was monitored by the number of associated ribosomes separated by sucrose gradient (M, monosome; P, polysome). The heatmap shows the codon frequency within the 9-nt random sequence in monosome (M) and polysome (P) fractions.
(D) A bar graph (right) shows the mean M/P ratio averaged across all positions in (C).
(E) The up panel shows the schematic of a massively parallel reporter assay, where a random 9-nt sequence was replaced before the uORF stop codon UAG. The uORF translation was monitored by the number of associated ribosomes separated by sucrose gradient (M, monosome; P, polysome). The heatmap shows the codon frequency within the 9-nt random sequence in monosome (M) and polysome (P) fractions.
(F) A bar graph shows the HiBiT signals in HEK293-K^b^ cells transfected with mRNA reporters bearing C-rich or GA-rich sequence before the GFP stop codon UAG. Error bars, mean ± s.e.m. *n* = 3 biological replicates. ***P* < 0.01 by unpaired two-tailed t-test.
(G) A bar graph shows the HiBiT signals in HEK293-K^b^ cells transfected with mRNA reporters bearing C-rich or GA-rich sequence before the GFP stop codon UAG. The downstream HiBiT sequence was inserted into different reading frames. Error bars, mean ± s.e.m. *n* = 3 biological replicates. ***P* ≤ 0.01 by unpaired two-tailed *t*-test.
(H) A bar graph shows the HiBiT signals in HEK293-K^b^ cells transfected with mRNA reporters bearing C-rich or GA-rich sequence before the GFP stop codon UAG. Error bars, mean ± s.e.m. *n* = 3 biological replicates. **P* < 0.05; ***P* ≤ 0.01; ****P* < 0.001 by unpaired two-tailed t-test for two groups and ordinary one-way ANOVA test followed by Dunnett's multiple comparisons for more than two groups.

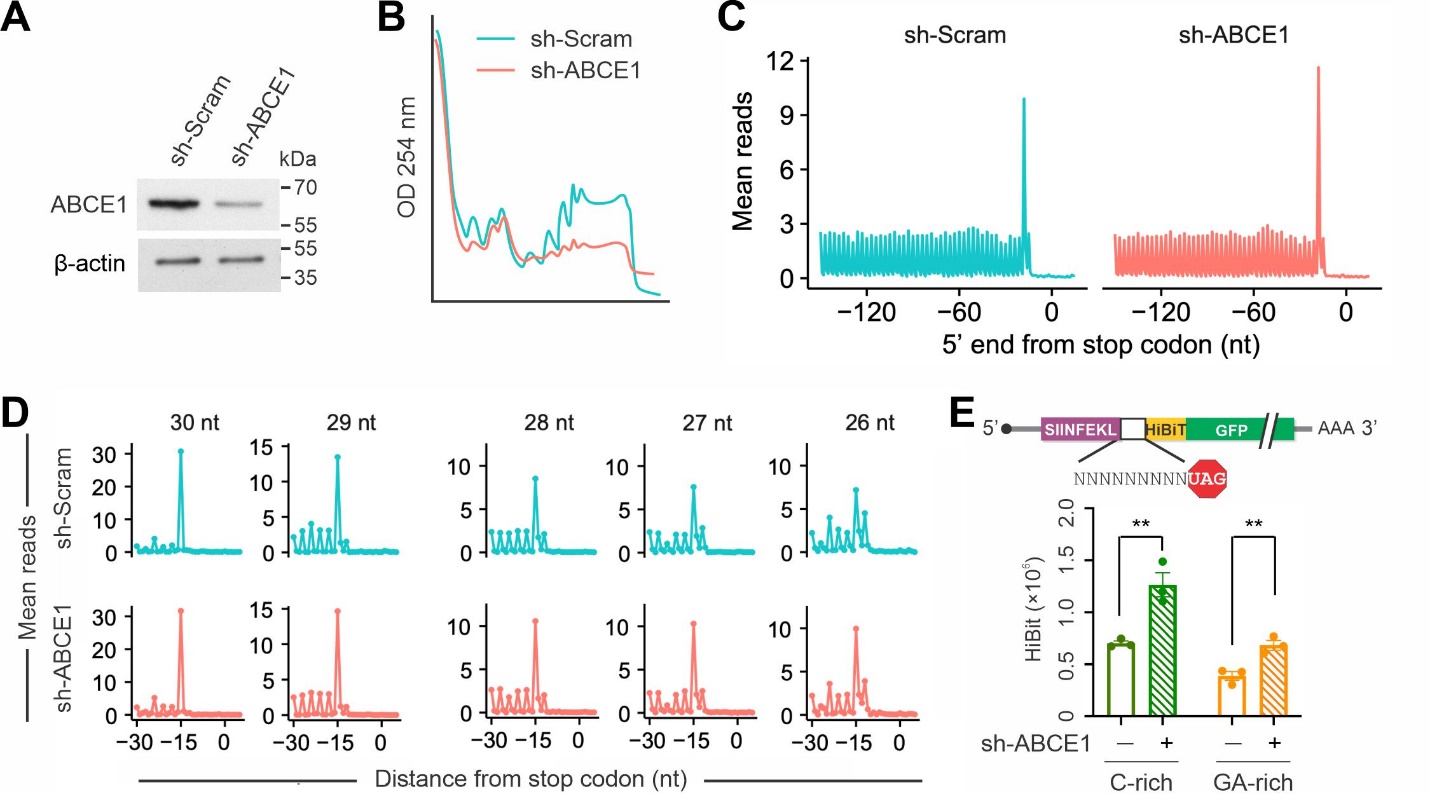

**Figure S4. ABCE1 modulates termination pausing independent of the mRNA sequence context.**

(A) Western blots of HEK293-K^b^ cells with or without ABCE1 knockdown.
(B) Comparison of polysome profiles of cells with or without ABCE1 knockdown.
(C) Aggregation plots show ribosome density around stop codons in cells with or without ABCE1 knockdown.
(D) Aggregation plots of mean Ribo-seq reads around the stop codon in cells with or without ABCE1 knockdown. The reads were stratified by the length followed by mapping using the 5’ end of footprint reads.
(E) A bar graph shows the HiBiT signals in HEK293-K^b^ cells with or without ABCE1 knockdown after transfection with mRNA reporters bearing C-rich or GA-rich sequence before the GFP stop codon UAG. Error bars, mean ± s.e.m. *n* = 3 biological replicates. ***P* < 0.01 by unpaired two-tailed t-test.

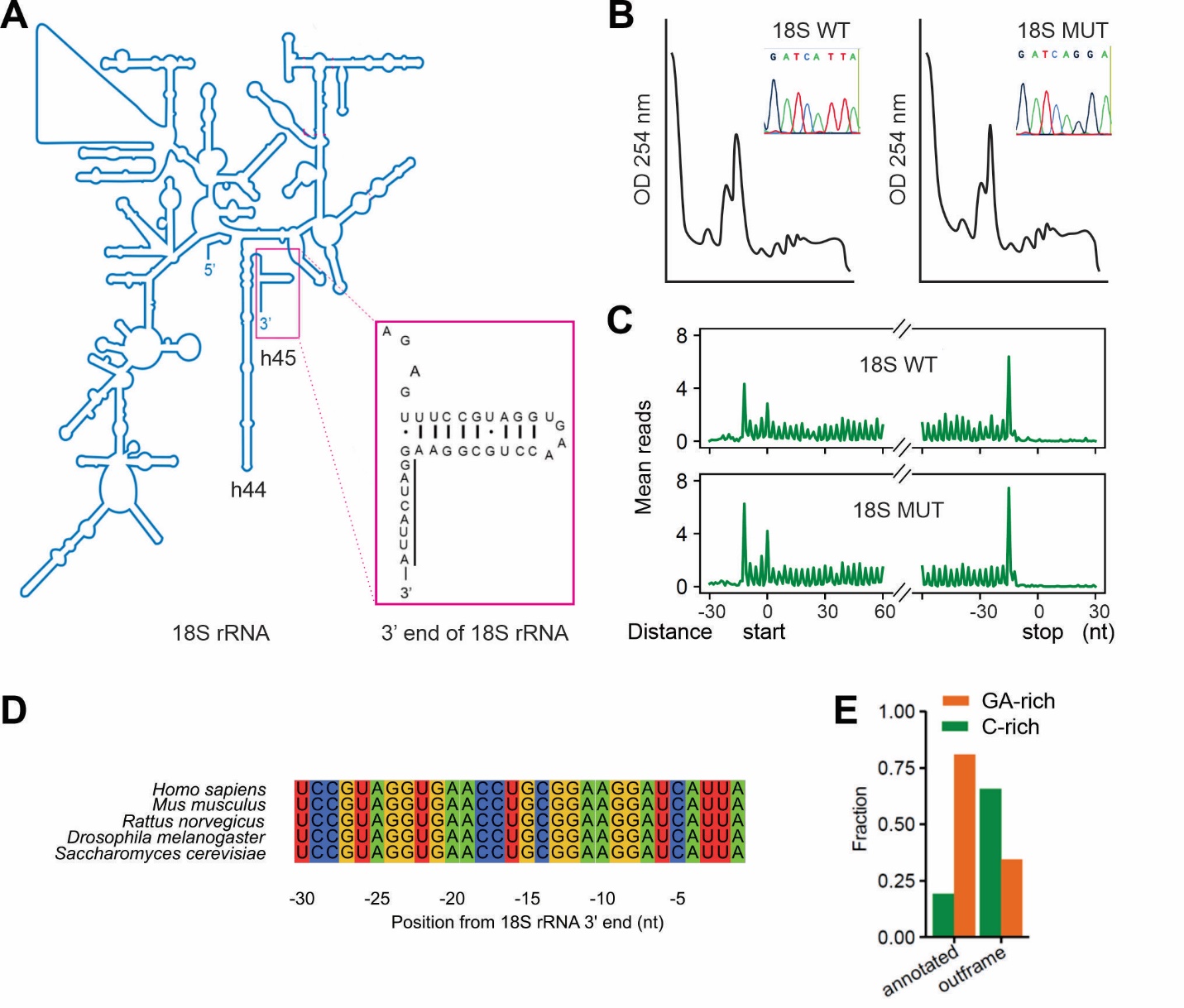

**Figure S5. The putative role of 3’ terminus of 18S rRNA in termination pausing.**

(A) A schematic shows the secondary structure of 18S rRNA with the 3’ end sequence highlighted.
(B) Comparison of polysome profiles of cells expressing the 18S rRNA WT or mutant. The inserts depict the nucleotide sequence of 18S rRNA WT or mutant.
(C) HEK293 cells were transfected with plasmids encoding 18S rRNA WT or mutant followed by Ribo-seq. Aggregation plots show the distribution of mean reads across the transcriptome aligned at start and stop codons.
(D) Sequence alignment of the 3’ end of 18S rRNA across multiple species.
(E) Relative frequency of stop codons preceded with GA-rich or C-rich seqeunces in human genome. Note the different fractions between annotated and out-of-frame stop codons.

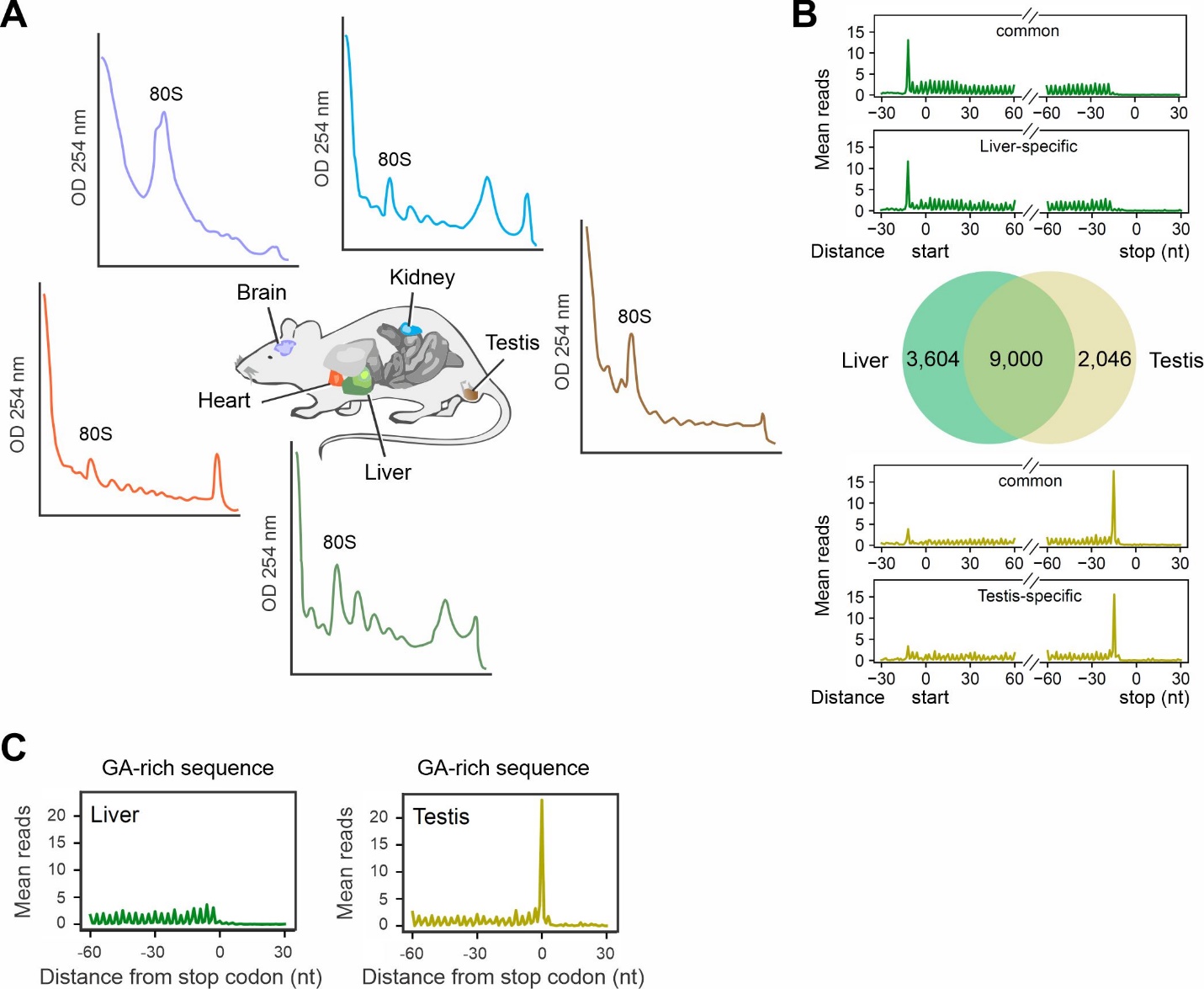

**Figure S6. Tissue specificity of termination pausing.**

(A) Different mouse tissues were collected followed by polysome profiling on sucrose gradients.
(B) Mouse liver and testis were subjected to Ribo-seq. Aggregation plots show the distribution of mean reads across the common or tissue-specific transcripts aligned at start and stop codons.
(C) Mouse liver and testis were subjected to Ribo-seq. Aggregation plots show the mean reads around stop codons of mRNAs with the GA sequence motif.

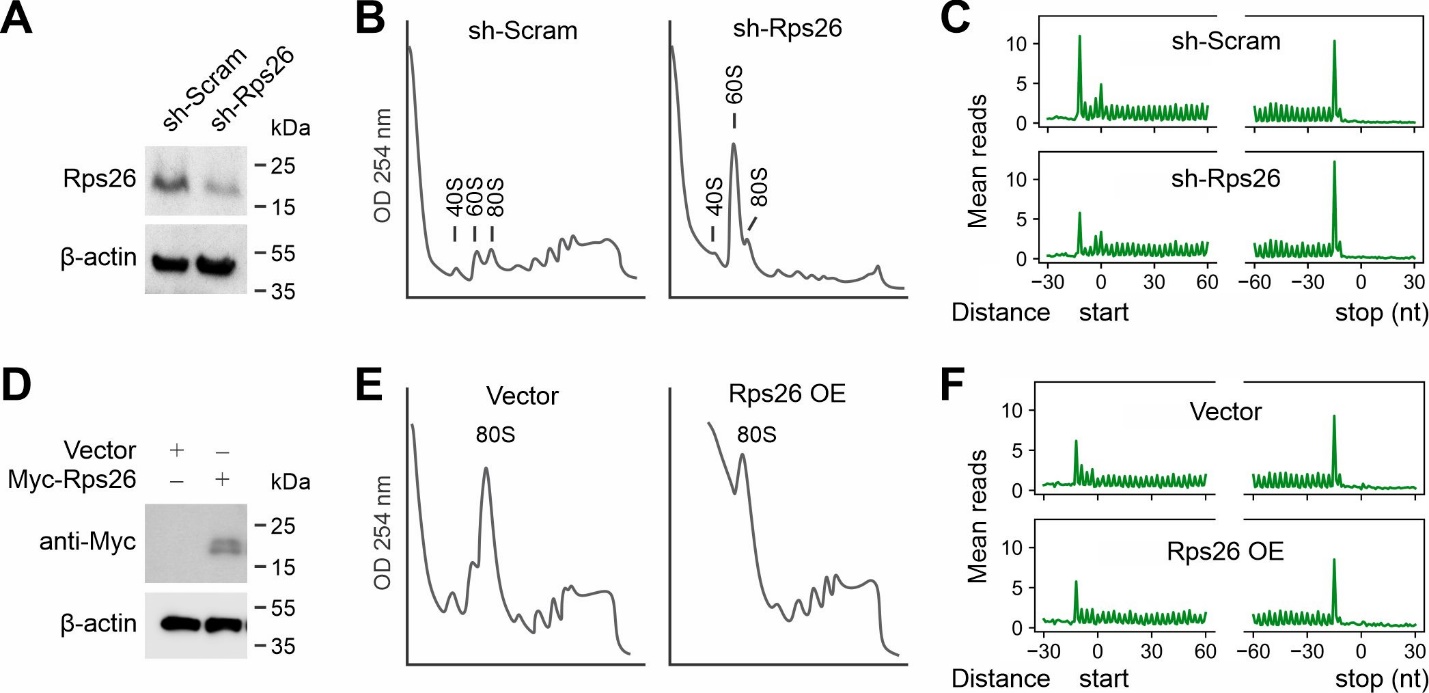

**Figure S7. The role of Rps26 in termination pausing.**

(A) Western blots of HEK293 cells with or without Rps26 knockdown.
(B) Comparison of polysome profiles of cells with or without Rps26 knockdown.
(C) HEK293 cells with or without Rps26 knockdown were subjected to Ribo-seq. Aggregation plots show the distribution of mean reads across the transcriptome aligned at start and stop codons.
(D) Western blots of HEK293 cells with or without Rps26 overexpression.
(E) Comparison of polysome profiles of cells with or without Rps26 overexpression.
(F) HEK293 cells with or without Rps26 overexpression were subjected to Ribo-seq. Aggregation plots show the distribution of mean reads across the transcriptome aligned at start and stop codons.
